# Epiphytic cryptogam communities differ little despite strong variation in coastal fog exposure

**DOI:** 10.64898/2026.10.16.759761

**Authors:** Abbey Schedler, Katrina McCollough, Frey Rogers, Kaylee Macomb, Michael J. Song, Jenna T.B. Ekwealor

## Abstract

Because mosses and lichens obtain water directly from their surroundings, coastal fog is expected to strongly influence epiphytic cryptogam communities during the long summer droughts of Mediterranean-climate ecosystems. We examined epiphytic mosses and lichens on coyote brush (*Baccharis pilularis*) across coastal and inland transects on San Bruno Mountain, California, where marine-layer exposure produces a sharp local microclimate gradient. We surveyed coyote brush availability, sampled standardized branches, monitored temperature and relative humidity, and quantified cryptogam occurrence, diversity, morphology, and reproduction. Morphological and chemical identifications were supplemented with molecular markers, while bacterial 16S sequencing was used to characterize cryptogam-associated microbiomes across host taxa and, for the dominant moss *Pulvigera pringlei*, across transects. Coastal microsites were cooler and more humid, whereas inland microsites experienced substantially greater daytime atmospheric drying demand. Despite this pronounced environmental contrast, cryptogam abundance, reproduction, and visible community composition differed little between transects, although branch-level genus diversity was modestly higher on the coastal transect. Macrolichen lobe or branch length and cryptogam patch area were greater inland in pooled comparisons, but these differences weakened after accounting for taxonomic composition. Bacterial communities differed strongly among cryptogam host taxa, but the *P. pringlei* microbiome showed no detectable difference in diversity or composition between coastal and inland transects. The broad similarity of cryptogam communities across a strong fog gradient is consistent with buffering of epiphytic cryptogams against fine-scale variation in marine-layer exposure. For the predominantly green-algal lichens in this community, the ability to absorb water vapor from humid air may reduce dependence on direct fog wetting, while host-associated bacterial communities may be structured more strongly by host identity than by local microclimate. Together, our results show that epiphytic cryptogam communities and microbiomes can remain remarkably similar across pronounced microclimatic variation despite modest morphological shifts.

## Introduction

Water availability limits biological activity in seasonally dry ecosystems, but its ecological effects depend on when and under what conditions organisms can use that water. During rainless periods, fog, dew, and humid air can provide direct hydration and reduce atmospheric water loss, extending opportunities for physiological activity (Fischer et al., 2009; Vasey et al., 2012; Emery, 2016). Fog can influence terrestrial plants directly, through the uptake of atmospheric water by leaves or other tissues, and indirectly, through fog drip and reduced evaporative demand (Potter, 2016; Emery, 2016; Emery et al., 2018). Greater atmospheric moisture might therefore be expected to increase growth in poikilohydric bryophytes and lichens, whose water status and physiological activity are closely coupled to environmental water availability (Proctor and Tuba, 2002; Borge and Ellis, 2024; Hanslin et al., 2001). However, the biological consequences of fog depend not only on water availability but also on the temperature and irradiance accompanying hydration. Persistent marine-layer conditions can cool surfaces and reduce light, potentially limiting carbon gain even while extending periods of physiological activity (Fischer et al., 2009; Borge and Ellis, 2024). Fog gradients may therefore produce strong environmental contrasts without necessarily producing equally strong differences in organismal performance or community composition.

Much of the research on non-rainfall water in coastal California has focused on vascular plants, including effects on plant water status, foliar uptake, and community composition (Vasey et al., 2012, 2014; Emery, 2016; Emery et al., 2018); broader reviews of fog–ecosystem interactions likewise focus largely on vascular plants (Weathers et al., 2020). Much less is known about how mosses and lichens interact with coastal fog and other forms of atmospheric moisture. These cryptogams are photosynthetic poikilohydric organisms: their internal water content rises and falls with external moisture, photosynthesis declines as tissues dry, and physiological activity resumes after rehydration (Proctor et al., 2007; Kranner et al., 2008; Stanton et al., 2023). This dependence on external water may be especially consequential for epiphytic mosses and lichens, which do not acquire water from soil through vascular roots and instead depend on water intercepted from the atmosphere and substrate surfaces (Gauslaa, 2014; Slate et al., 2024). Rain provides episodic liquid-water inputs, whereas fog, dew, and humid air can provide additional or repeated hydration during otherwise rainless periods (Gauslaa, 2014; Slate et al., 2024).

Dependence on atmospheric moisture does not necessarily imply dependence on direct fog wetting. Fog consists of suspended liquid water droplets, whereas humid air can hydrate thalli without visible wetting. Many green-algal lichens containing *Trebouxia* can resume photosynthetic activity after desiccation through hydration from humid air alone, whereas cyanolichens generally require liquid water for photosynthetic activation (Phinney et al., 2018, 2019; Lange et al., 1986). For lichens in particular, vapor pressure deficit is especially relevant because water content and physiological activity respond strongly to atmospheric drying demand (Stanton et al., 2023). For bryophytes, tolerance of desiccation and rehydration are shaped not only by whether tissues dry, but by the rate of drying, water content reached, duration of dry periods, and conditions during rehydration (Proctor et al., 2007; Stark, 2017). Cryptogam responses to coastal fog may therefore reflect both the amount and form of atmospheric water and the environmental conditions accompanying hydration.

Microclimate can affect cryptogam communities at several levels that need not respond at the same rate. Strong environmental filtering could alter which taxa establish or persist, whereas established individuals may instead respond through changes in size, morphology, or reproductive output. Moreover, lichen morphology is closely related to water acquisition and retention, with thin or highly branched thalli absorbing water vapor rapidly and thicker forms retaining liquid water for longer periods (Gasulla et al., 2021; Phinney et al., 2018). Fine-scale fog gradients have also been associated with differences in lichen growth form, orientation, abundance, and species composition (Stanton and Horn, 2013; Stanton, 2015). Cryptogam hosts can selectively structure their associated microbial communities, potentially reducing the influence of external environmental variation (Grube et al., 2009; Wísniewski et al., 2025; Mawarda et al., 2026). Measuring community composition, organismal morphology, reproduction, and host-associated microbiomes together therefore allows us to distinguish broad community turnover from subtler biological responses within an otherwise persistent community.

Cryptogam communities contain important biological variation that is not apparent from morphology alone. This is especially true for lichens, whose visible thalli arise from associations among fungal, algal, cyanobacterial, and bacterial partners rather than from a single organism (Grube et al., 2009; Grimm et al., 2021). Molecular markers can supplement morphological and chemical identification when visible characters alone do not resolve cryptogam identity. Cryptogams also host diverse bacterial communities that may vary with host identity, thallus or shoot morphology, and the surrounding environment (Grube et al., 2009; Grimm et al., 2021; Chen and Nelson, 2022). These associations can be functionally important. In mosses, associated bacteria can contribute to nutrient acquisition and nitrogen fixation and can influence host growth and development, including through production of plant hormones (Alvarenga and Rousk, 2022; Hornschuh et al., 2006; Schauer and Kutschera, 2011). We therefore combined visible community surveys and molecular identification with bacterial microbiome sequencing to characterize both cryptogam community structure and host-associated bacterial communities.

San Bruno Mountain, on the central California coast, provides a natural setting in which to examine cryptogam responses to fine-scale variation in marine-layer exposure (Fig. 1a–c). The mountain lies between the Pacific Ocean and San Francisco Bay, and its main ridge separates a coast-facing slope that experiences persistent summer fog from a warmer inland-facing slope. Although San Bruno Mountain supports extensive coastal scrub and chaparral vegetation within a highly urbanized landscape, its bryophyte and lichen communities remain poorly documented relative to its vascular flora and animal communities (McClintock and Fahy, 1968; Allshouse and Nelson, 2022). Coyote brush (*Baccharis pilularis* DC.) is widespread in coastal California shrublands, occurs across both maritime and more inland environments, and provides an abundant woody substrate for epiphytic mosses and lichens (Fig. 1d–e). It can also intercept and directly use fog water (Emery, 2016; Emery et al., 2018).

**Figure 1.**
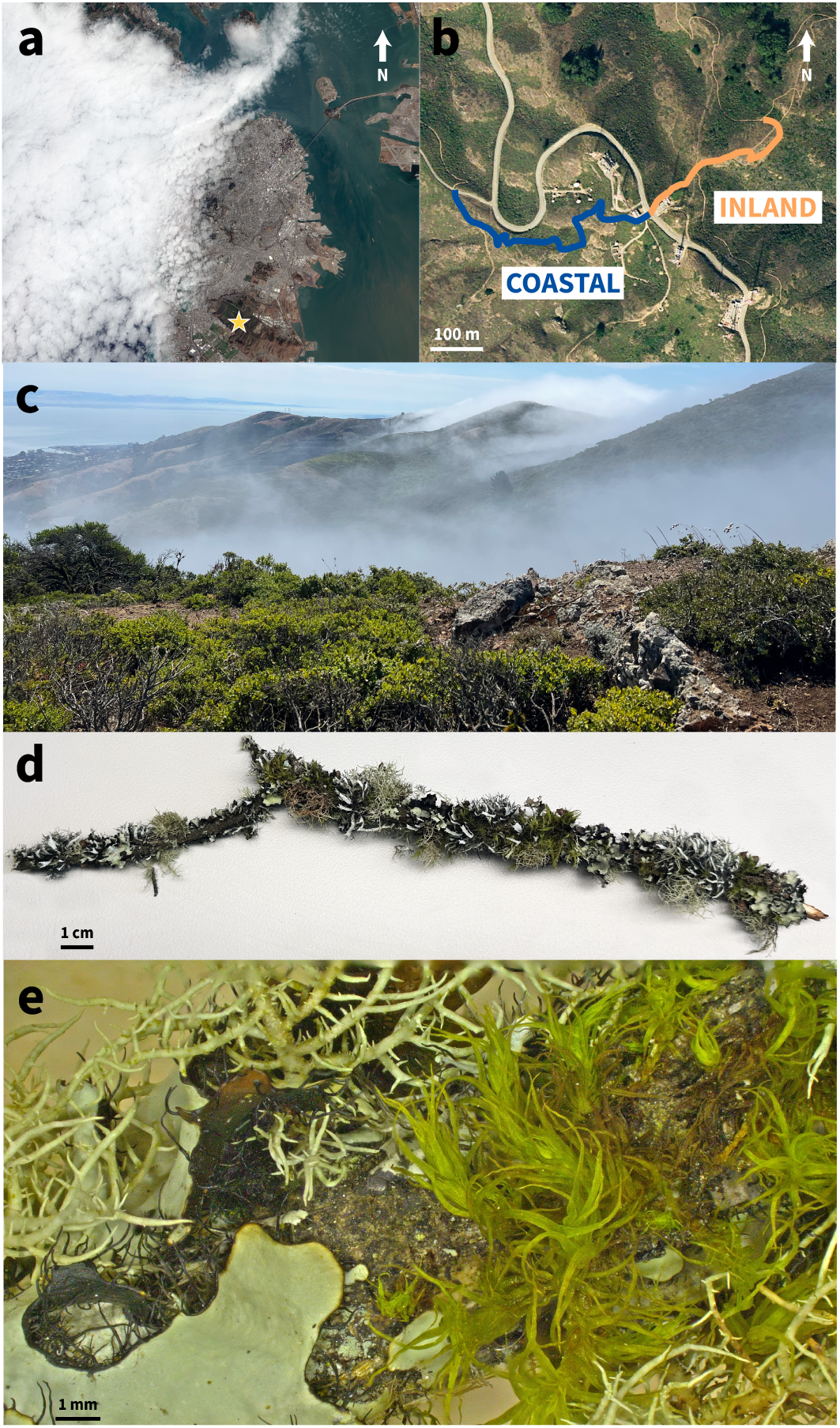
Study site, sampling design, and epiphyte community at San Bruno Mountain, California. (a) Regional satellite image showing coastal fog across the San Francisco Peninsula; the star indicates San Bruno Mountain. Base image: NASA Earth Observatory. (b) Coastal and inland sampling transects on San Bruno Mountain. Base image: USGS EarthExplorer. (c) Marine-layer fog moving across coastal scrub habitat at San Bruno Mountain. Photo: Jenna Ekwealor. (d) Representative epiphyte-covered branch used for community and morphological measurements; scale bar = 1 cm. (e) Foliose lichen *Parmotrema perlatum* and moss *Pulvigera pringlei* growing on a sampled branch; scale bar = 1 mm.

Here, we examined whether epiphytic cryptogam communities on *B. pilularis* differed across the coastal–inland microclimatic gradient on San Bruno Mountain. We asked whether abundance, diversity, morphology, reproduction, and community composition differed between transects, whether cryptogam host identity structured associated bacterial communities, and whether the microbiome of the dominant moss, *Pulvigera pringlei*, varied with transect or moss morphology. We initially expected the cooler and more humid coastal transect to support greater cryptogam abundance, diversity, size, and reproductive activity because atmospheric moisture can extend physiological activity during the summer dry season. Two alternatives, however, could weaken or reverse this response. Cryptogams capable of using water vapor may remain hydrated under humid conditions even without direct fog deposition, reducing dependence on fog wetting itself, while persistent coastal fog may also reduce temperature and irradiance during hydrated periods and thereby limit carbon gain.

## Materials and methods

### Study area and sampling design

San Bruno Mountain State & County Park is located on the central California coast, immediately south of San Francisco and between the Pacific Ocean and San Francisco Bay (Fig. 1a). The mountain is surrounded by extensive urban development but supports coastal scrub and chaparral vegetation along its main ridge. The coastal scrub includes *Anaphalis margaritacea*, *Eriophyllum staechadifolium*, *Diplacus aurantiacus*, *Frangula californica*, and *Artemisia californica*, among other species. Marine-layer fog frequently reaches the mountain during the summer dry season, producing a strong microclimate contrast across the ridge (Fig. 1c). The sampled ridge runs approximately northwest to southeast. We established a pair of coastal and inland transects descending from a shared high point near the ridge at 37.68887° N, 122.43703° W (Fig. 1b). The coastal transect extended southeast from Radio Road along the Summit Loop Trail toward Battery 59 and ranged from approximately 314 to 365 m elevation. The inland transect extended northwest from Radio Road along the Summit Loop Trail toward its intersection with the Connector and Dairy Ravine trails and ranged from approximately 338 to 366 m elevation. We collected all branch and cryptogam samples in July and August 2024. Field sampling was conducted under San Mateo County Parks scientific research permit SP-2024-07-SBM.

Coyote brush (*Baccharis pilularis* DC.) occurred at the site in both prostrate and upright growth forms. We sampled only upright shrubs, restricting sampling to a single host species and growth form to reduce variation in substrate identity and host architecture across transects. We selected 20 shrubs, including 10 along each transect. Shrubs were selected to be similar in size and growth form across transects.

### Host availability survey

We conducted a supplementary survey in July of 2025 to determine whether the availability of upright *B. pilularis* differed between transects. We surveyed seven points along the coastal transect and 12 points along the inland transect. Survey points were spaced at approximately 100-m intervals along each transect. At each point, we counted all upright *B. pilularis* at least 1 m tall whose central trunk fell within a 3-m-radius circle centered on the survey point. We also recorded the height of the tallest upright shrub, the presence or absence of the prostrate growth form, elevation, latitude, and longitude. These measurements were used to evaluate whether differences between transects in epiphyte occurrence or potential landscape-scale abundance could reflect differences in the availability or size of suitable host shrubs.

### Microclimate monitoring and processing

We deployed 14 DS1923 Hygrochron temperature and relative-humidity loggers (iButton; Maxim Integrated, San Jose, California, USA), with seven placed along each transect. We attached each logger to a branch at a similar height on a *B. pilularis* shrub similar in size and growth form to the shrubs used for epiphyte sampling. Loggers were launched and downloaded using OneWireViewer and were configured to record temperature and relative humidity at 15-min intervals. Loggers were deployed on 20 June 2025. At this sampling interval, the loggers could store approximately 43 d of observations. We first retrieved and relaunched available loggers on 6 August 2025, 47 d after deployment, and completed a second retrieval on 26 September 2025, after an additional 51 d. Logger records varied in duration because some units reached memory capacity before retrieval, failed during deployment, or could not be relaunched after the first retrieval. We retained all available observations and summarized each logger over the period for which it was active; sample sizes therefore varied among dates and summary periods.

We imported and processed temperature and relative-humidity records in R 4.5.2 using tidyverse and lubridate. We assigned each logger to the coastal or inland transect from its serial number, parsed timestamps in the America/Los Angeles time zone, and constrained relative humidity values to 0–100%. We corrected a documented timestamp offset for one logger and removed a short interval when that logger was not recording field conditions during retrieval. We classified each 15-min observation as daytime or nighttime using date-specific sunrise and sunset times calculated for San Bruno Mountain in the America/Los Angeles time zone. These classifications were retained when aggregating observations into hourly summaries, allowing hours spanning sunrise or sunset to contain separate daytime and nighttime means.

We calculated saturation vapor pressure following Buck (1981) as

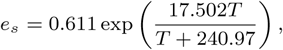

where *T* is temperature in degrees Celsius. We calculated actual vapor pressure as *e_a_* = *e_s_*(*RH/*100), where *RH* is relative humidity, and calculated vapor pressure deficit (VPD) as *e_s_ − e_a_*, in kPa.

### Host branch collection and measurements

From each selected shrub, we collected one branch approximately 1 cm in diameter and 29 cm long from approximately 1.25 m above the ground. Branches were transported to the laboratory in unsealed plastic bags and stored air-dried under cool, dark conditions until processing, which took place over the next two years.

For each branch, we designated two opposing surfaces as side A and side B and retained this orientation for all subsequent measurements. During laboratory processing, we measured the length and diameter of each branch segment and counted lateral branches. Lateral branches were removed during surveys; when two branch segments from the same sample were processed separately, their measured lengths were summed for branch-level analyses. We used measured branch length to standardize comparisons of patch counts, patch area, and reproductive structures between transects and to evaluate sampling effort in the diversity analysis.

### Cryptogam surveys

We surveyed visible epiphytic cryptogams on each sampled branch and treated each spatially discrete, measurable patch as an individual observation. For each patch, we recorded taxonomic identity, morphology, and reproductive structures.

We distinguished patch-based macroepiphyte records from microepiphyte records operationally. Macroepiphyte patches were moss or lichen patches large enough to distinguish and measure as discrete patches. Smaller, crustose, microscopic, or more dispersed epiphytes that could not be measured as discrete patches were retained only as branch-level presence–absence records. Very small leafy liverworts occurred occasionally near branch bases, often growing over lichen thalli, and were treated as microepiphytes under this criterion.

For mosses, we defined a measurable patch as a spatially discrete aggregation containing at least two shoots. Isolated shoots were excluded from patch counts, including occasional long shoots growing within lichen patches. We photographed each moss patch with a metric ruler positioned in the same plane as the patch and measured projected patch area from photographs using ImageJ.JS, the browser-based version of ImageJ (Abràmoff et al., 2004). For each moss patch, we selected the longest shoot, rehydrated it with deionized water, and measured shoot length. Smaller shoots were measured with a Leica Ivesta 3 stereomicroscope with an integrated camera, and larger shoots were measured with a metric ruler when they exceeded the microscope field of view. We measured the length of three fully hydrated leaves collected from the middle portion of the shoot and used their mean as the patch-level leaf-length value. When possible, leaves were taken from the same shoot measured for shoot length; otherwise, leaves were removed from the second-longest shoot without removing the entire shoot. We recorded the presence and number of sporophytes in each moss patch as a measure of sexual reproduction.

For macrolichens, we recorded branch side and position along the branch. Patch position was recorded using a string marked at 1-cm intervals; for example, a record spanning 3–5 cm indicated that the patch began 3 cm from the branch origin and extended to 5 cm. We measured patch length, width, and height, and measured the maximum lobe or branch length and width for each patch. Foliose lichen lobes were measured as individual projecting lobes, whereas fruticose lichen branches were measured as individual filamentous branches. Measurements were made after lightly moistening thalli with water to reduce breakage during handling. We recorded the presence or absence of sexual and asexual reproductive structures for each macrolichen patch. Macrolichens were identified morphologically to genus or species when possible using standard keys and chemical spot tests when needed.

Because multiple patches could occur on the same branch, host branch was retained as the sampling unit for branch-level analyses of patch abundance, diversity, and reproductive structure density.

### Molecular identification of cryptogams and symbiotic partners

#### Sample preparation and direct PCR

We prepared lichen and moss tissues for direct PCR using a low-biomass cryptogam workflow adapted from Jung et al. (2024). For each lichen sample, we washed target thallus fragments with deionized water until hydrated. Under a dissecting microscope, we removed small fragments using sterile forceps and cleaned them of visible contaminants. When necessary, we adjusted tissue fragment size or mechanically disrupted samples using a pestle or pipette tip to improve amplification success.

We prepared crude DNA lysates using Platinum Direct PCR Universal Master Mix (Invitrogen, Thermo Fisher Scientific, Waltham, Massachusetts, USA; catalog no. A44647100). For each preparation, we added 20 *µ*L of lysis buffer, 0.6 *µ*L of proteinase K, and prepared tissue to a 1.5 mL microcentrifuge tube. We heated tubes at 98 °C for 1 min; cyanolichen samples were heated for 2 min. For samples requiring further optimization, we diluted crude lysates 1:10 before PCR.

#### Marker amplification

We amplified marker regions directly from crude moss and lichen lysates using the same Platinum Direct PCR Universal Master Mix. For lichen samples, we amplified mycobiont rDNA using ITS1-F/LR3, chlorobiont rDNA using Al1500af/LR3, and cyanobiont 16S rRNA using CYA361f/CYA785r. For moss samples, we amplified ITS1 and ITS2 separately. For bacterial 16S reactions used in the cross-host microbiome analysis, we amplified nearly full-length 16S rRNA using the degenerate S-D-Bact-0008-c-S-20/S-D-Bact-1492-a-A-22 primer pair with Oxford Nanopore-compatible 5*^′^* anchor sequences. Primer and PNA sequences, approximate amplicon sizes, and sources are provided in Supplementary Table S4.

Each 20 *µ*L PCR reaction contained 10 *µ*L master mix, 1 *µ*L of premixed forward and reverse primers, and 2 *µ*L DNA lysate. For bacterial 16S reactions, we additionally included anti-plastid (pPNA-S; PP01-25) and anti-mitochondrial (mPNA-S; MP01-25) peptide nucleic acid clamps (PNA Bio), using approximately 1 *µ*L of each clamp to reduce amplification of host organellar 16S sequences (Lundberg et al., 2013). We adjusted the volume of deionized water to maintain a final reaction volume of 20 *µ*L.

For fungal ITS, algal ITS, and bacterial 16S reactions, we used an initial denaturation at 94 °C for 2 min, followed by 35 cycles of 94 °C for 15 s, 60 °C for 15 s, and 68 °C for 20 s, followed by storage at 4 °C. For moss ITS reactions, ITS1 and ITS2 were amplified separately using an initial denaturation at 94 °C for 1 min, followed by 30 cycles of 94 °C for 1 min, primer-specific annealing for 1 min, and 72 °C for 1.5 min, with a final extension at 72 °C for 5 min. The annealing temperature was 52 °C for ITS1 and 59 °C for ITS2.

We assessed PCR success by gel electrophoresis and cleaned successful PCR products using the NucleoSpin Gel and PCR Clean-up Kit (Macherey–Nagel, Düren, Germany). For cleanup, we diluted 15 *µ*L PCR product with 35 *µ*L deionized water before following the kit protocol.

#### Nanopore sequencing and taxonomic assignment

We pooled cleaned PCR products for nanopore sequencing. We prepared libraries using the Oxford Nanopore Rapid Barcoding Kit 24 V14 (SQK-RBK114.24) and sequenced them on a single R10 Flongle Flow Cell using a MinION Mk1D device.

We trimmed raw nanopore reads using cutadapt v5.2 (Martin, 2011), removing 76 bases from the 5*^′^* end of each read (-u 76). We imported trimmed reads into Geneious 2025.2 and generated taxonomic assignments using NCBI BLAST (Altschul et al., 1990) through the Geneious platform. We reviewed BLAST assignments and retained identifications at the finest reliable taxonomic level. We used fungal ITS, algal ITS, moss ITS, and cyanobacterial markers to confirm or refine morphological and chemical identifications of cryptogams and their symbiotic partners.

#### Pulvigera pringlei bacterial microbiome DNA extraction, amplification, and sequencing

For the dedicated *P. pringlei* bacterial microbiome analysis, we sampled one *P. pringlei* patch from each of the branches collected in 2024, after approximately two years of dry storage, and extracted DNA in June–July 2026. One sample failed to yield usable sequencing data and was excluded from microbiome analyses, leaving 19 samples. For each patch, we measured the longest shoot, photographed the patch with a metric ruler for area measurement in ImageJ (Abràmoff et al., 2004), and recorded moss tissue mass. We measured patch area as the area containing moss growth, including portions where moss was intertwined with lichen.

We placed moss tissue in a 1.5 mL tube with 400 *µ*L ATL buffer. When an entire patch could not be accommodated, we used the maximum amount of tissue that fit in a single tube. We incubated samples overnight at 4 °C, shook them for 5 min at maximum speed, and briefly centrifuged them. We removed 250 *µ*L of buffer and followed the ZymoBIOMICS DNA Miniprep Kit protocol for liquid samples. Because DNA concentrations were consistently low, we concentrated extracts on a heat block for 25 min at 65 °C.

Test PCRs with 16S primers confirmed bacterial amplification, and we submitted samples to Plasmidsaurus for 16S amplicon sequencing and taxonomic profiling. Plasmidsaurus amplified full-length 16S, sequenced libraries using Oxford Nanopore long-read sequencing, and generated taxonomic profiles with its standard Microbiome 16S workflow. Briefly, Plasmidsaurus classified quality-filtered reads against rrnDB v5.6 and the NCBI Targeted Loci database using emu v3.5.1 (Curry et al., 2022) and reported taxa comprising at least 0.1% of each microbial community.

## Statistical analyses

We conducted all data processing and statistical analyses in R 4.5.2. Analysis code will be made available at https://github.com/ abbeykate/SBM_fog. Unless otherwise specified, tests were two-sided. Permutation-based community analyses used 999 permutations. We used tidyverse packages for data preparation, lubridate for date-time handling, vegan for diversity, dissimilarity, PERMANOVA, and dispersion analyses, ape for principal coordinates analysis, and ggplot2, cowplot, and patchwork for figures.

### Microclimate

We first averaged observations within logger, metric, transect, and hour. We then averaged logger-level hourly means within each transect so that each logger received equal weight, and we calculated standard errors among logger means. For time-series comparisons, we retained only hours with measurements from both transects. We calculated inland-minus-coastal differences for temperature and VPD and coastal-minus-inland differences for relative humidity. We propagated uncertainty in transect differences as the square root of the sum of squared transect standard errors and broke plotted lines across gaps longer than 3 h.

We classified observations as daytime or nighttime using date-specific sunrise and sunset times for all temperature, relative-humidity, and VPD summaries. To quantify the midday temperature contrast, we averaged hourly inland-minus-coastal temperature differences between 10:00 and 15:59 local time within each date and tested whether the mean daily difference departed from zero using a two-sided one-sample *t*-test. We also used two-sided one-sample *t*-tests to evaluate mean daily daylight differences in relative humidity and VPD.

We also calculated daily daytime VPD separately for each transect. We first averaged VPD across loggers within each hour and then gave each hourly mean equal weight in the daily mean. We retained transect-days with at least eight daytime hours and calculated standard errors across hourly means.

For each date, we calculated the proportion of matched daylight hours with vapor pressure deficit (VPD) exceeding 1.5 kPa separately for each transect. We compared paired daily proportions using a two-sided exact sign test, excluding days with identical proportions.

Because many green-algal lichens can use atmospheric water vapor under sufficiently humid conditions (Lange et al., 1986; Phinney et al., 2018, 2019), we also summarized periods of very low VPD. For this exploratory summary, we classified VPD *<* 0.3 kPa as very low; this threshold was used to identify highly humid conditions rather than to represent a physiological cutoff. Using the hourly transect-level summaries, we calculated the proportion of observations below this threshold overall and separately during solar-defined daytime and nighttime. For time-matched hours, we additionally quantified how often both transects, only the coastal transect, only the inland transect, or neither transect fell below 0.3 kPa. We also summarized the duration of consecutive low-VPD periods, treating observations as consecutive only when separated by exactly 1 h. Because hourly observations were temporally autocorrelated, these threshold summaries were interpreted primarily as descriptive rather than as independent physiological tests.

### Cryptogam abundance and diversity

We treated branch as the replicate for branch-level comparisons and individual patches as observations for patch-level trait comparisons. We quantified sampling effort as measured branch length. For community summaries, we combined macrolichen patch measurements, moss patch measurements, and branch-level microepiphyte records. We treated microepiphytes as presence– absence records, whereas macrolichens and mosses were treated as counted patch records.

We calculated branch-level diversity after combining macrolichens, mosses, and microepiphytes. We summed records within each genus and branch, calculated observed genus richness, and calculated Shannon entropy from genus-level counts. We exponentiated Shannon entropy to express Hill diversity of order 1 as the effective number of genera. To assess whether unequal sampled branch lengths influenced the comparison, we fitted a linear model with Hill diversity of order 1 as the response and transect and measured branch length as covariates.

We compared total epiphyte patch density between transects with a negative-binomial generalized linear model using patch count as the response, transect as the predictor, and the log of measured branch length as an offset. We summarized transect-level patch area as total positive patch area divided by total measured branch length.

We compared positive-valued patch area, macrolichen maximum lobe or branch length, moss shoot length, and branch-level Hill diversity between inland and coastal transects with two-sided Wilcoxon rank-sum tests using the large-sample approximation. For these tests, we evaluated macrolichen maximum lobe or branch length, moss shoot length, and patch area at the patch level, whereas we evaluated diversity at the branch level. For Wilcoxon tests displayed in figure panels, we reported the Wilcoxon statistic, Hodges–Lehmann location shift, and associated *P* value.

Because macrolichen genera differ substantially in characteristic thallus size, we conducted a sensitivity analysis to determine whether the pooled inland–coastal difference in maximum lobe or branch length reflected differences in taxonomic composition between transects. We fitted a linear mixed-effects model to log-transformed maximum lobe or branch length, with transect and genus as fixed effects and host branch as a random intercept to account for multiple lichen patches measured on the same branch. We evaluated the contribution of transect by comparing the full model with a reduced model lacking transect using a likelihood-ratio test. As a robustness check, we repeated the analysis using only genera represented by at least five observations on each transect. We also summarized inland–coastal differences within genera represented by at least five observations on each transect. To assess whether the taxonomic composition of macrolichen observations contributing to the size analysis differed between transects, we compared genus-by-transect counts using a Fisher’s exact test with simulated *P*-values.

Because patch area was also analyzed across multiple cryptogam taxa, we conducted an analogous sensitivity analysis to determine whether the pooled inland–coastal difference in positive patch area reflected taxonomic composition. We fitted a linear mixed-effects model to log-transformed patch area, with transect and genus as fixed effects and host branch as a random intercept. We evaluated the contribution of transect by comparing the full model with a reduced model lacking transect using a likelihood-ratio test. As a robustness check, we repeated the analysis using only genera represented by at least five observations on each transect. We also summarized inland–coastal differences within genera represented by at least five observations on each transect. We compared genus-by-transect counts using a Fisher’s exact test with simulated *P*-values.

### Reproduction and host availability

We analyzed macrolichen reproductive output at the branch level after standardizing by total macrolichen patch area. For each branch, we calculated sexual reproductive patch density as the number of macrolichen patches with sexual structures per cm^2^ of macrolichen patch area and asexual reproductive patch density as the number of macrolichen patches with asexual structures per cm^2^ of macrolichen patch area. We analyzed moss reproductive output as the number of capsules per cm^2^ of moss patch area. We compared each reproductive response between transects with a two-sided Wilcoxon rank-sum test using the large-sample approximation.

For the ancillary coyote-brush survey, we summarized maximum shrub height and number of upright shrubs meeting the survey size criterion by transect. We assessed normality within each transect with Shapiro–Wilk tests. We compared maximum shrub height between transects with a two-sided Welch two-sample *t*-test. Because shrub counts were not normally distributed within at least one transect, we compared counts with a Wilcoxon rank-sum test.

### Visible community composition

We described visible cryptogam community composition from patch counts for each genus–species combination within each transect. To account for differences in total sampled branch length between transects, we normalized patch abundance by dividing each taxon’s summed patch count by total sampled branch length within the transect and rescaling to the mean total sampled branch length across transects. For Fig. 4, we displayed normalized abundances as stacked bars to show overall community composition (Fig. 4a) and as paired taxon-specific values to show the direction and magnitude of inland–coastal differences (Fig. 4b). In exploratory genus-level screens, we compared branch-level occurrence of each genus between transects with Fisher’s exact tests. For quantitatively counted genera, we also modeled branch-level patch counts with negative-binomial generalized linear models containing transect and an offset for sampled branch length. We adjusted *P*-values in these exploratory genus-level screens using the Benjamini–Hochberg false-discovery-rate procedure.

### Bacterial alpha and beta diversity

For microbiome analyses across cryptogam hosts, we analyzed bacterial 16S records only. We used fungal ITS and moss ITS records for host or symbiont identification and excluded them from bacterial microbiome richness and composition analyses. We identified each cryptogam replicate by lysate ID plus host taxon because some lysate labels were shared by different host cryptogams. We standardized the diluted rerun lysate 11-2.D to 11-2 and did not treat it as an independent biological lysate.

For cross-host richness, we retained curated bacterial 16S records identified to genus and calculated richness as the number of distinct bacterial genera recovered for each cryptogam replicate. For cross-host composition, we counted repeated assignments to the same genus within a replicate as recovered sequences and converted them to within-replicate relative sequence abundance. We calculated Bray– Curtis dissimilarities from genus-level relative sequence abundance and represented relationships among samples with two-dimensional principal coordinates analysis using an additive correction for non-Euclidean distances. We tested the effect of host taxon on bacterial community composition with PERMANOVA and tested homogeneity of multivariate dispersion with betadisper and a permutation test. To assess whether host-associated differences were driven by individual samples, we repeated PERMANOVA after sequentially excluding each lysate (leave-one-out sensitivity analysis).

For the dedicated *P. pringlei* bacterial microbiome analysis, we imported Plasmidsaurus relative-abundance tables and linked samples to branch and transect metadata. We retained bacterial taxa with positive reported relative abundance and summarized communities at the genus level. Before diversity and composition analyses, we removed unresolved taxon labels. We did not rarefy the data.

We calculated genus-level Hill diversity of order 1 as the exponential of Shannon entropy and compared it between transects with a two-sided Wilcoxon rank-sum test. Genus-level Bray–Curtis dissimilarities were calculated from relative-abundance profiles and represented with two-dimensional principal coordinates analysis using an additive correction. We tested the effect of transect on *P. pringlei* bacterial community composition with PERMANOVA and assessed homogeneity of multivariate dispersion with betadisper and a 999-permutation test. For descriptive composition plots, genera were ranked by mean relative abundance across samples; the 10 most abundant genera were displayed separately; all remaining genera were pooled as “Other”; and displayed relative abundances were renormalized to sum to one within each sample. although inland microsites experienced substantially greater daytime drying demand, both environments regularly experienced prolonged periods of very low VPD.

### Host shoot length and bacterial community structure

For each *P. pringlei* microbiome sample, we joined moss morphology measurements from the extraction metadata to genus-level microbiome diversity by branch. We tested associations between moss shoot length and genus-level Hill diversity of order 1 with Spearman rank correlation using the large-sample approximation. We used the same procedure for moss patch area and genus-level Hill diversity. For visualization, we displayed points by transect; inference was based on the Spearman tests.

## Results

### A strong microclimate contrast separated coastal and inland transects

Inland microsites were warmer and experienced greater atmospheric drying demand than coastal microsites (Fig. 2). High-VPD (*>* 1.5 kPa) conditions occurred more frequently on the inland transect. Of 86 days with matched daylight observations, 14 had a greater proportion of high-VPD hours inland, none had a greater proportion coastal, and 72 showed no difference (two-sided exact sign test, *P* = 0.00012). Midday temperatures were significantly higher inland, with a mean daily inland-minus-coastal difference of 1.05 °C (95% CI: 0.86–1.25 °C; two-sided one-sample *t*-test, *P* = 2.84 *×* 10*^−^*^17^; *n* = 86 days; Fig. 2a). Temperature differences were greatest during the day, although inland microsites frequently remained warmer overnight. The largest temperature differences occurred during August and September.

**Figure 2.**
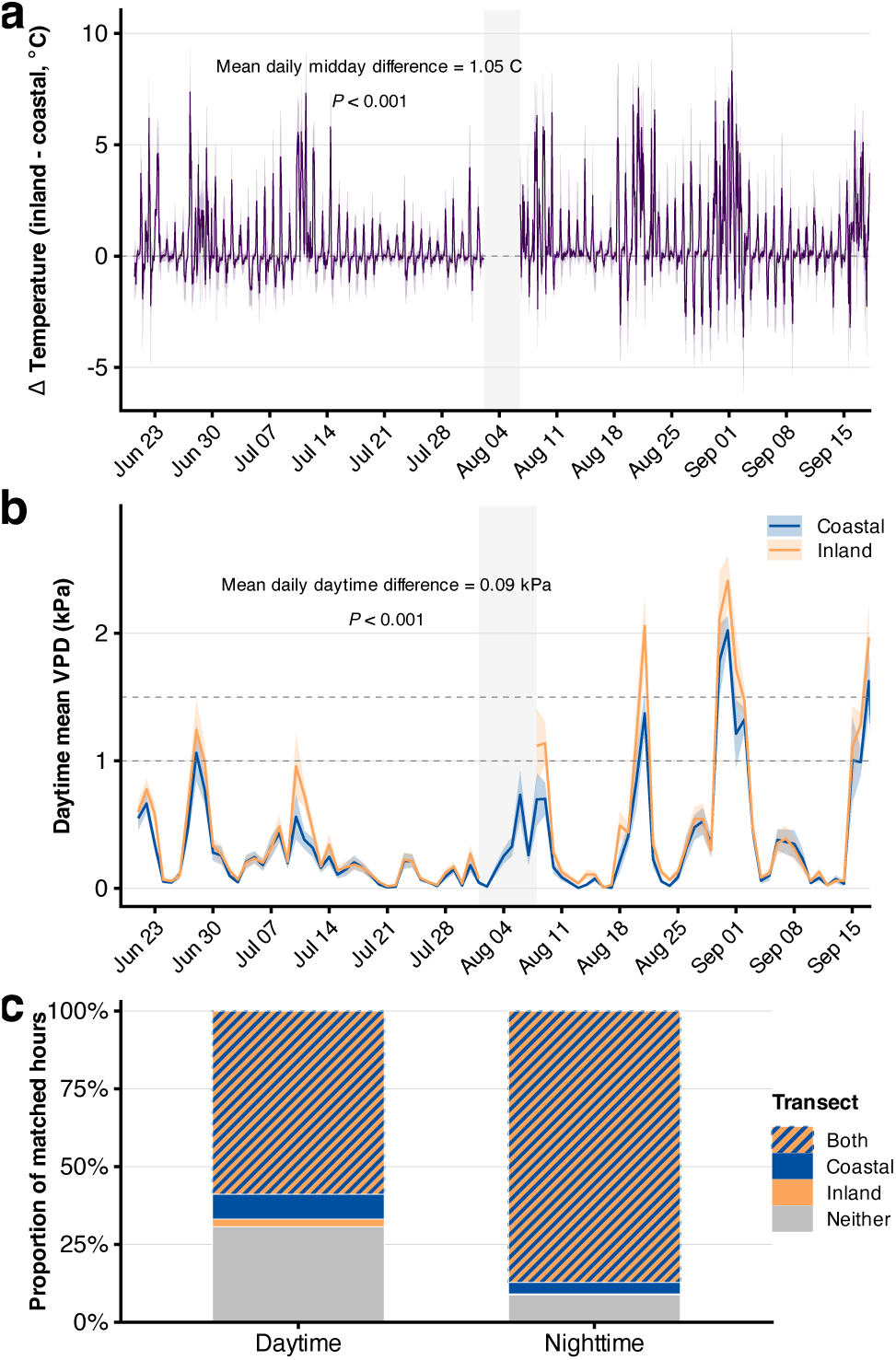
Microclimate differences between coastal and inland transects. (a) Hourly inland-minus-coastal temperature differences with *±*1 standard error ribbons; positive values indicate warmer conditions inland. Gray shading indicates data gaps. The annotation reports the mean daily midday (10:00– 15:59 local time) difference and its two-sided one-sample *t*-test *P*-value. (b) Daily daytime mean vapor pressure deficit (VPD) for coastal and inland transects, with *±*1 standard error ribbons calculated across daytime hourly means. Daytime was defined using local sunrise and sunset. Gray shading indicates gaps in the inland record, and dashed horizontal lines mark 1.0 and 1.5 kPa. The annotation reports the mean daily inland-minus-coastal daytime VPD difference and its two-sided one-sample *t*-test *P*-value. (c) Proportion of time-matched daytime and nighttime hours in which VPD was below 0.3 kPa on both transects, only the coastal transect, only the inland transect, or neither transect. The 0.3-kPa value was used to identify highly humid conditions for this exploratory summary and does not represent a physiological cutoff.

Despite this greater daytime atmospheric drying demand inland, very low-VPD conditions occurred frequently on both transects. VPD was below 0.3 kPa during 75.2% of coastal and 71.5% of inland hourly observations. These conditions were especially common at night, occurring during 89.7% of coastal and 87.5% of inland nighttime observations. Among 945 time-matched nighttime hours, both transects were simultaneously below 0.3 kPa during 824 hours (87.2%). Very low-VPD conditions also frequently extended into daylight hours: both transects were simultaneously below 0.3 kPa during 743 of 1,261 matched daytime hours (58.9%; Fig. 2c). Low-VPD conditions were generally sustained rather than isolated, with a median consecutive duration of 17 h on both transects. Thus,

### Host availability did not differ between transects

Neither the abundance of suitable coyote brush hosts nor maximum shrub height differed significantly between transects (Wilcoxon rank-sum test for shrub counts: *W* = 39, *P* = 0.8253; Welch two-sample *t*-test for shrub height: *P* = 0.40).

### Cryptogam communities were broadly similar across transects despite modest morphological differences

Cryptogam abundance differed little between transects, whereas branch-level Hill diversity was modestly higher on coastal branches. Total epiphyte patch density was similar between the coastal and inland transects, but branch-level Hill diversity was higher on the coastal transect (median Inland = 5.11, Coastal = 5.73; Hodges– Lehmann shift, Inland minus Coastal =-0.83; *W* = 21, *P* = 0.031; Fig. 3e). This difference remained evident after accounting for measured branch length in a sensitivity analysis (coastal effect = 0.93 effective genera, 95% CI = 0.19–1.68, *P* = 0.017), whereas branch length itself was not associated with diversity (*P* = 0.381).

**Figure 3.**
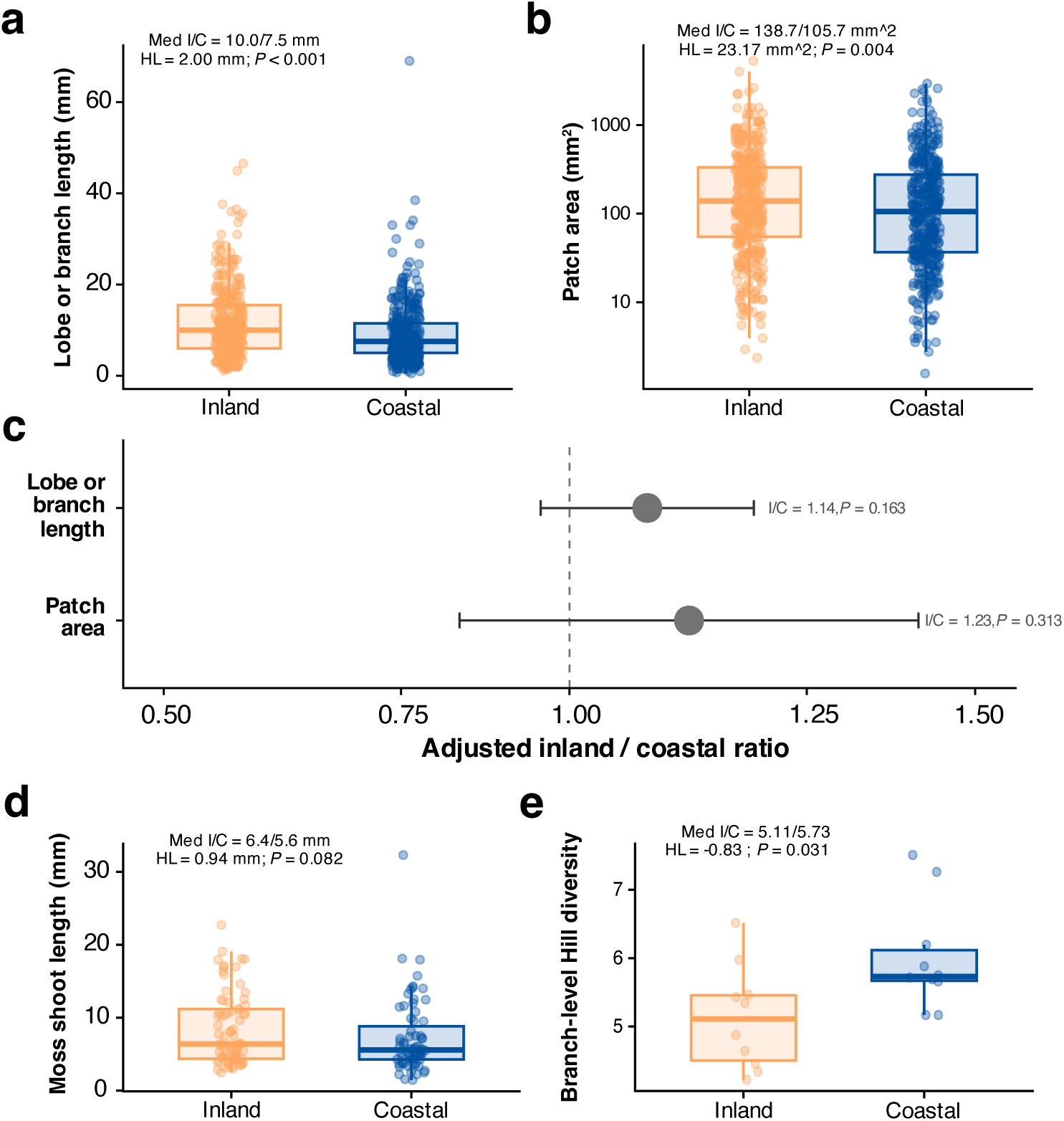
Cryptogam morphology and diversity responses across inland and coastal transects. (a) Pooled macrolichen maximum lobe or branch length and (b) pooled positive cryptogam patch area, shown as raw observations and boxplots by transect; annotations give inland/coastal medians, Hodges–Lehmann location shifts, and Wilcoxon rank-sum *P*-values. (c) Genus-and branch-adjusted inland/coastal geometric-mean ratios for maximum lobe or branch length and patch area from linear mixed-effects models; points show estimated ratios, horizontal lines show 95% confidence intervals, and the dashed vertical line marks no difference between transects (ratio = 1). (d) Moss shoot length and (e) branch-level Hill diversity (q = 1) by transect, shown as raw observations and boxplots with inland/coastal medians, Hodges–Lehmann location shifts, and Wilcoxon rank-sum *P*-values.

**Figure 4.**
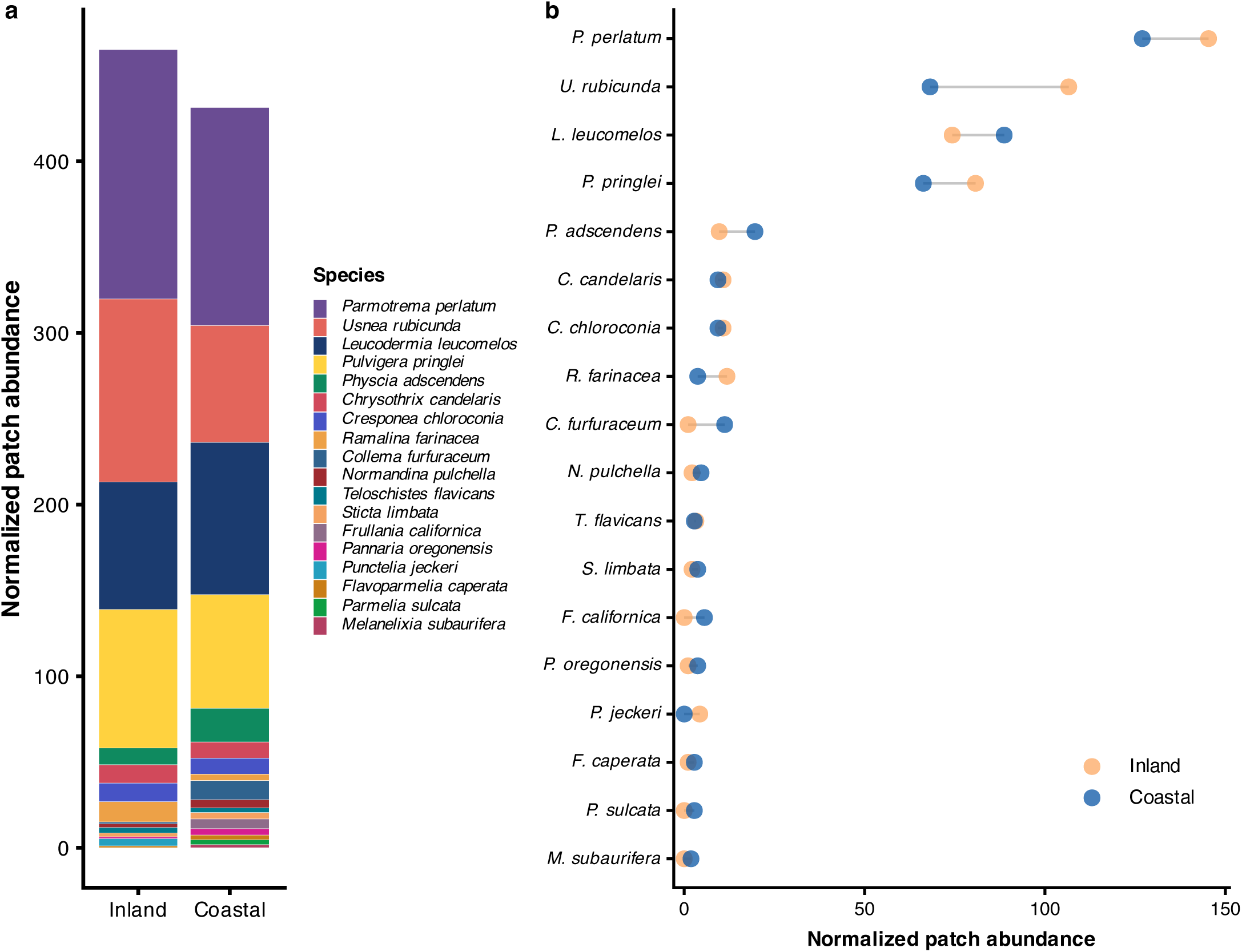
Cryptogam community composition and abundance across inland and coastal transects. (a) Stacked bars show normalized patch abundance by taxon on the inland and coastal transects. (b) Paired points show the same normalized abundance for each taxon; gray lines connect inland and coastal values for the same taxon. Abundance was normalized for differences in sampled branch length by dividing each taxon’s total patch count by total sampled branch length within its transect and rescaling to the mean total sampled branch length across transects. Taxa are shown in the same order in both panels to facilitate comparison.

Morphological differences were more apparent in pooled comparisons but were modest after accounting for taxonomic composition. Maximum macrolichen lobe or branch length was greater inland in the pooled comparison (median 10.0 vs 7.5 mm; Hodges–Lehmann difference = 2.0 mm, *P* = 7.3 *×* 10*^−^*^7^; Fig. 3a), but the adjusted model estimated a 14.2% inland increase, which was not statistically significant (95% CI for the inland/coastal ratio: 0.95–1.37; likelihood-ratio *P* = 0.163; Fig. 3c). Positive patch area was also greater inland in the pooled comparison (median 138.7 vs 105.7 mm^2^; Hodges–Lehmann difference = 23.17 mm^2^, *P* = 0.00368; Fig. 3b), but the adjusted model estimated a 22.7% inland increase, which was not statistically significant (95% CI for the inland/coastal ratio: 0.83–1.82; likelihood-ratio *P* = 0.313; Fig. 3c). Within-taxon directions were mixed for both responses, indicating that the pooled differences did not reflect a uniform within-taxon response to transect (Supplementary Figs. S1 and S2). Moss shoot length did not differ detectably between transects (Fig. 3d). Two-sided Wilcoxon rank-sum tests likewise detected no transect differences in macrolichen sexual reproductive patch density (*W* = 55, *P* = 0.690), asexual reproductive patch density (*W* = 54, *P* = 0.791), or moss capsule density (*W* = 54, *P* = 0.772).

### Visible cryptogam community composition differed little between transects

Visible cryptogam community composition was broadly similar between the coastal and inland transects (Fig. 4a). The same dominant moss and macrolichen taxa occurred on both transects. Normalized patch abundance showed modest taxon-specific differences, but the overall dominance structure remained similar (Fig. 4b). No individual cryptogam genus showed a significant difference in occurrence or abundance between transects after correcting for multiple comparisons. Cyanolichen patches were more frequent on the coastal transect (20 vs. 4 measured patches), although cyanolichens were uncommon overall.

### Cryptogam host identity structured bacterial communities

Bacterial community profiles were generated for 24 samples representing 11 cryptogam host taxa, enabling comparisons of bacterial diversity and community composition among hosts. Bacterial genus richness varied among cryptogam host taxa, with individual samples containing 1–31 genera and host-level means ranging from 1 to 18 genera (Fig. 5a). Bacterial community composition differed significantly among host taxa (PERMANOVA: *R*^2^ = 0.54, *P* = 0.002; Fig. 5b), whereas multivariate dispersion did not differ among host taxa (dispersion test: *P* = 0.106). Host-associated differences were robust to the exclusion of individual lysates, with leave-one-out PERMANOVA results ranging from *R*^2^ = 0.514 to 0.574 and *P* = 0.001 to 0.018.

**Figure 5.**
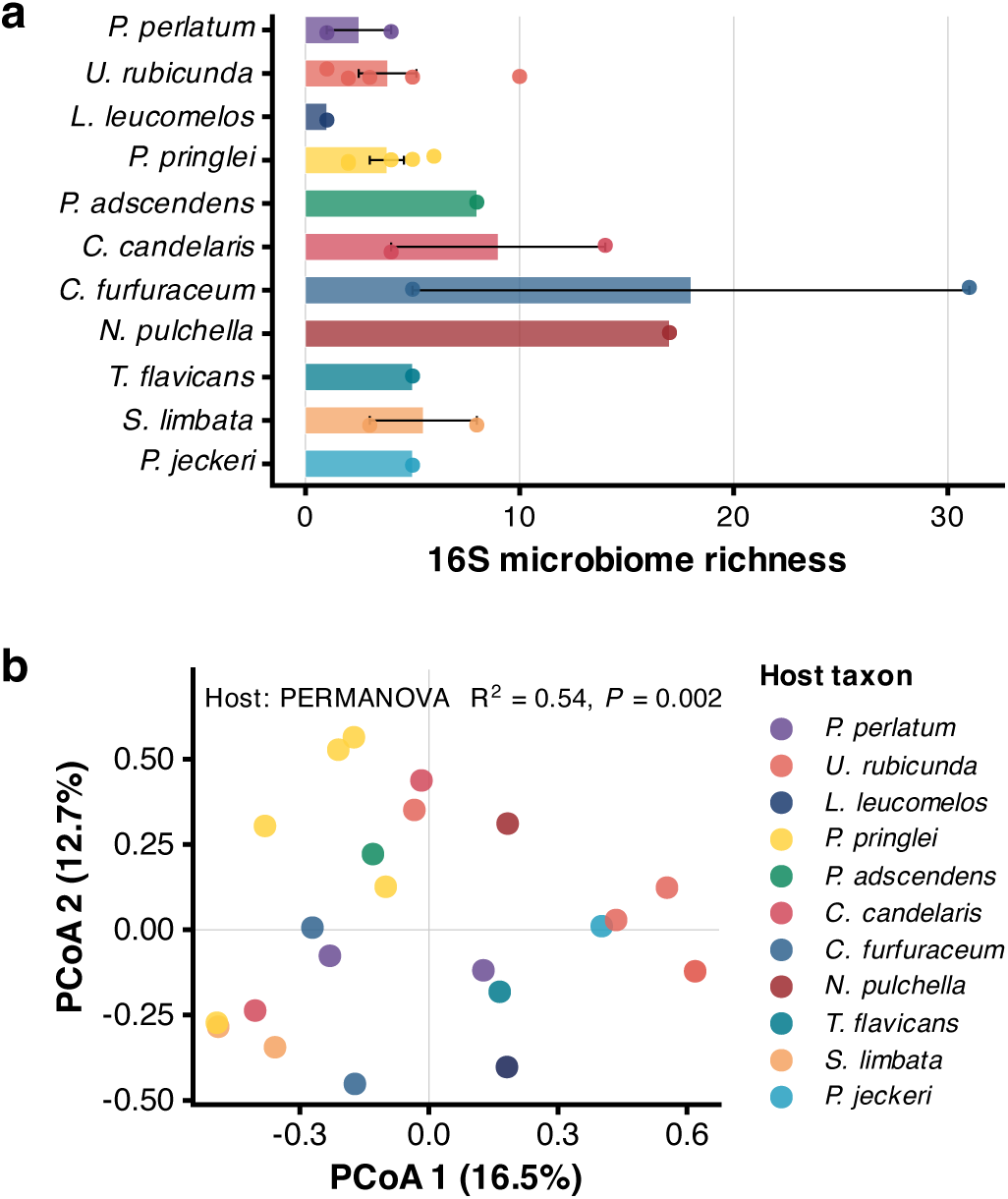
Bacterial community diversity and composition across cryptogam hosts. (a) Mean observed bacterial genus richness across lysates for each cryptogam host taxon; points show individual lysates, bars show means, and error bars indicate *±*1 standard error of the mean. (b) Principal coordinates analysis of bacterial community composition based on Bray– Curtis dissimilarities; points are colored by host taxon. Bacterial community composition differed among host taxa (PERMANOVA: *R*^2^ = 0.54, *P* = 0.002), whereas multivariate dispersion did not differ among hosts (*P* = 0.106).

### The bacterial microbiome of *Pulvigera pringlei* differed little across the microclimatic gradient

Genus-level bacterial alpha diversity did not differ significantly between inland and coastal *P. pringlei* samples (Wilcoxon rank-sum test: *W* = 66, *P* = 0.094; Fig. S3a), nor did bacterial community composition (PERMANOVA: *R*^2^ = 0.042, *P* = 0.670; Fig. S3b). Multivariate dispersion was also similar between transects (dispersion test: *P* = 0.469), and the same dominant bacterial genera occurred in both environments (Fig. S4). Bacterial genus diversity was unrelated to moss shoot length or patch area (Fig. S5). A summary of microbiome statistical tests is provided in Supplementary Table S3.

## Discussion

Despite a pronounced microclimatic contrast across San Bruno Mountain, epiphytic cryptogam communities differed surprisingly little between coastal and inland transects. The inland transect was warmer and experienced substantially greater daytime atmospheric drying demand than the coastal transect (Fig. 2). Yet cryptogam abundance, reproductive investment, visible community composition, and the bacterial microbiome of the dominant moss *Pulvigera pringlei* were broadly similar across the ridge (Fig. 4; Supplementary Figs. S3 and S4). The similar availability of upright *B. pilularis* on the two transects further suggests that comparable epiphyte abundance was not masking a large transect-scale difference driven by host availability. Branch-level cryptogam genus diversity was modestly higher on the coastal side, while pooled lobe length and patch area were greater inland; both morphological differences weakened after accounting for taxonomic composition (Fig. 3). For lobe length in particular, this weakening indicates that the pooled inland difference was driven in part by greater representation of larger-lobed taxa rather than by a consistent inland increase within taxa. Taken together, these patterns suggest that the microclimatic gradient affected some aspects of local community structure and morphology without producing broad turnover in the epiphyte assemblage.

One explanation for the similarity between transects is the temporal distribution of atmospheric moisture. Both transects regularly experienced low VPD; both were simultaneously below 0.3 kPa during 87.2% of matched nighttime hours, and continuous low-VPD periods lasted a median of 17 h on each transect (Fig. 2c). Thus, although the inland transect experienced stronger daytime atmospheric drying demand, periods of higher VPD were repeatedly interrupted by sustained humid conditions. For poikilohydric cryptogams, this recurrence may be important because overnight hydration could allow tissues to begin each morning hydrated, while low-VPD conditions that persist into daylight may extend physiological activity after sunrise and, for some green-algal lichens, permit vapor hydration while light is available (Phinney et al., 2018, 2019; Borge and Ellis, 2024). Together, these observations suggest that the biological consequences of the coastal–inland contrast may depend not only on the total time available for hydration, but also on how frequently humid periods recur and how recently tissues were hydrated before atmospheric drying demand increases during the day. This temporal structure may have buffered cryptogams against the stronger daytime microclimatic contrast, helping to explain why community composition, reproduction, and host-associated microbiomes differed so little across the ridge. The modest increase in branch-level diversity coastal nevertheless suggests that this buffering was incomplete and that fine-scale community structure remained responsive to the microclimatic gradient (Fig. 3e).

The morphological differences we observed were subtler than the pooled comparisons initially suggested. Both maximum macrolichen lobe or branch length and positive patch area were greater inland when all taxa were analyzed together, but these effects weakened after accounting for taxonomic identity (Fig. 3a–c). Differences in the relative representation of larger-bodied taxa contributed to the pooled patterns, although they did not explain them completely. Within-taxon responses were heterogeneous, with some taxa tending to be larger inland and others showing little difference or the opposite direction (Supplementary Figs. S1 and S2). Such heterogeneous responses are plausible because lichen taxa differ substantially in thallus architecture and water relations, so the same change in humidity or drying demand need not produce the same morphological response across taxa (Gasulla et al., 2021; Phinney et al., 2018). The pooled inland differences therefore do not reflect a general tendency for cryptogams to grow larger inland.

Visible cryptogam community composition was notably conserved across the ridge (Fig. 4). In contrast, Stanton and Horn (2013) documented substantial changes in epiphytic lichen cover, community composition, and growth form across a coastal fog gradient in semiarid Chile. Their findings demonstrate that fog gradients can strongly structure epiphyte communities, making the limited turnover observed at San Bruno Mountain particularly noteworthy. Higher branch-level Hill diversity on the coastal transect (Fig. 3e), despite little genus-level turnover, indicates that the coastal difference was expressed primarily within individual branches rather than through replacement of the ridge-scale epiphyte assemblage. Coastal branches therefore supported a slightly richer or more even mix of genera, or both. Cyanolichens also showed a suggestive taxon-specific pattern, with 20 measured patches on the coastal transect and only four on the inland transect. Cyanolichens were too uncommon for strong inference from this contrast, but their greater representation on the coastal transect is qualitatively consistent with their stronger dependence on liquid water for photosynthetic activation relative to many green-algal lichens (Lange et al., 1986; Phinney et al., 2019). Broad similarity in community composition therefore coexisted with subtler responses in local diversity and in taxa that differ in how they acquire atmospheric water.

Cryptogam-associated bacterial communities showed clear host-associated structure. Bacterial community composition differed strongly among cryptogam host taxa (Fig. 5), whereas within *P. pringlei*, neither bacterial alpha diversity nor community composition differed significantly between coastal and inland samples, and the same dominant bacterial genera occurred on both transects (Supplementary Figs. S3 and S4). Bacterial diversity within *P. pringlei* was also unrelated to moss shoot length or patch area (Supplementary Fig. S5). The strong differentiation among host taxa is consistent with host-associated filtering playing an important role in structuring these bacterial communities, while the weak differentiation within *P. pringlei* indicates little microbiome response to the coastal–inland contrast.

The weak transect response of the *P. pringlei* microbiome is consistent with growing evidence that cryptogam-associated bacterial communities can remain similar across contrasting environments. In the epiphytic lichen *Evernia prunastri*, bacterial diversity and overall community composition remained similar between urban and natural environments, with most bacterial taxa shared between habitats (Mawarda et al., 2026). Comparable patterns have been reported in contribute to nutrient acquisition, particularly nitrogen fixation, and have been implicated in pathogen defense, carbon exchange, and regulation of host growth (Alvarenga and Rousk, 2022). Experimental work with *Funaria hygrometrica* has shown that epiphytic *Methylobacterium* can increase protonemal cell length and cell number, with auxin production implicated in the growth-promoting effect (Hornschuh et al., 2006). Earlier studies likewise showed that naturally associated bacteria can alter moss growth and gametophore development (Spiess et al., 1984). We did not measure microbial function in the present study, so these mechanisms remain hypotheses for *P. pringlei*; however, they provide plausible routes through which host-associated bacterial communities could influence cryptogam performance even when microbiome composition changes little across the microclimatic gradient.

More broadly, our results indicate that a strong microclimatic gradient did not translate into comparably strong biological turnover. Coastal and inland microsites on San Bruno Mountain differed strongly in temperature and atmospheric drying demand (Fig. 2), yet most components of the epiphytic cryptogam community remained similar (Figs. 3 and 4; Supplementary Fig. S3). The temporal distribution of atmospheric moisture may help explain this broad similarity: recurrent humid periods could repeatedly provide opportunities for hydration or slow water loss despite stronger daytime drying inland, while persistent coastal fog may also reduce temperature and irradiance during periods when carbon gain is possible. At the same time, modest differences in branch-level diversity and morphology show that the coastal–inland gradient was not biologically inconsequential. Further comparisons across liverworts. In *Riccia*, bacterial diversity and community composition were relatively insensitive to field environment, with a core bacterial community shared across field types and locations (Wicaksono et al., 2023), while the bacteriome of *Calasterella californica* retained similar alpha diversity and a prevalent core set of bacterial taxa across widely separated regions of California despite geographic and climatic variation in non-core taxa (González-Ramírez et al., 2026). The distinct thallus microbiome of *Riccia sorocarpa* relative to surrounding soil further suggests that cryptogam hosts can selectively structure their associated microbial communities (Wísniewski et al., 2025). Together with the strong differences we observed among cryptogam host taxa (Fig. 5), these studies suggest that host identity may structure cryptogam-associated bacterial communities more strongly than fine-scale environmental variation. However, our sample size limits the detection of subtle transect differences within *P. pringlei*, and the nonsignificant results do not establish equivalence between its coastal and inland bacterial communities. Because microbiome samples were obtained from branches stored dry for approximately two years, storage may also have influenced the recovered bacterial communities. Such host-associated structure could also have functional consequences for the cryptogam. Moss-associated bacteria can independent coastal–inland gradients will help establish how broadly these patterns apply. Predicting cryptogam responses to changing coastal fog regimes will also require distinguishing changes in visible fog frequency from changes in the humidity, temperature, and light conditions that ultimately determine physiological activity.

## Data availability

Processed data and analysis code supporting this study will be made available at https://github.com/abbeykate/SBM_fog. Raw sequence reads will be deposited in the NCBI Sequence Read Archive (SRA) prior to journal publication.

## Author contributions statement

J.T.B.E. conceived the study. J.T.B.E., K.A.M., F.R., and A.K.S. designed the experiments and analytical approaches. A.K.S., K.A.M., J.T.B.E., K.M., and F.R. performed fieldwork, laboratory work, and data collection. All authors contributed to data analysis and interpretation. All authors contributed to manuscript writing and approved the final manuscript.

## Acknowledgments

We thank San Mateo County Parks and California State Parks for permission to conduct fieldwork at San Bruno Mountain State & County Park.

## Supplementary Information

Supplementary Figures

**Figure S1.**
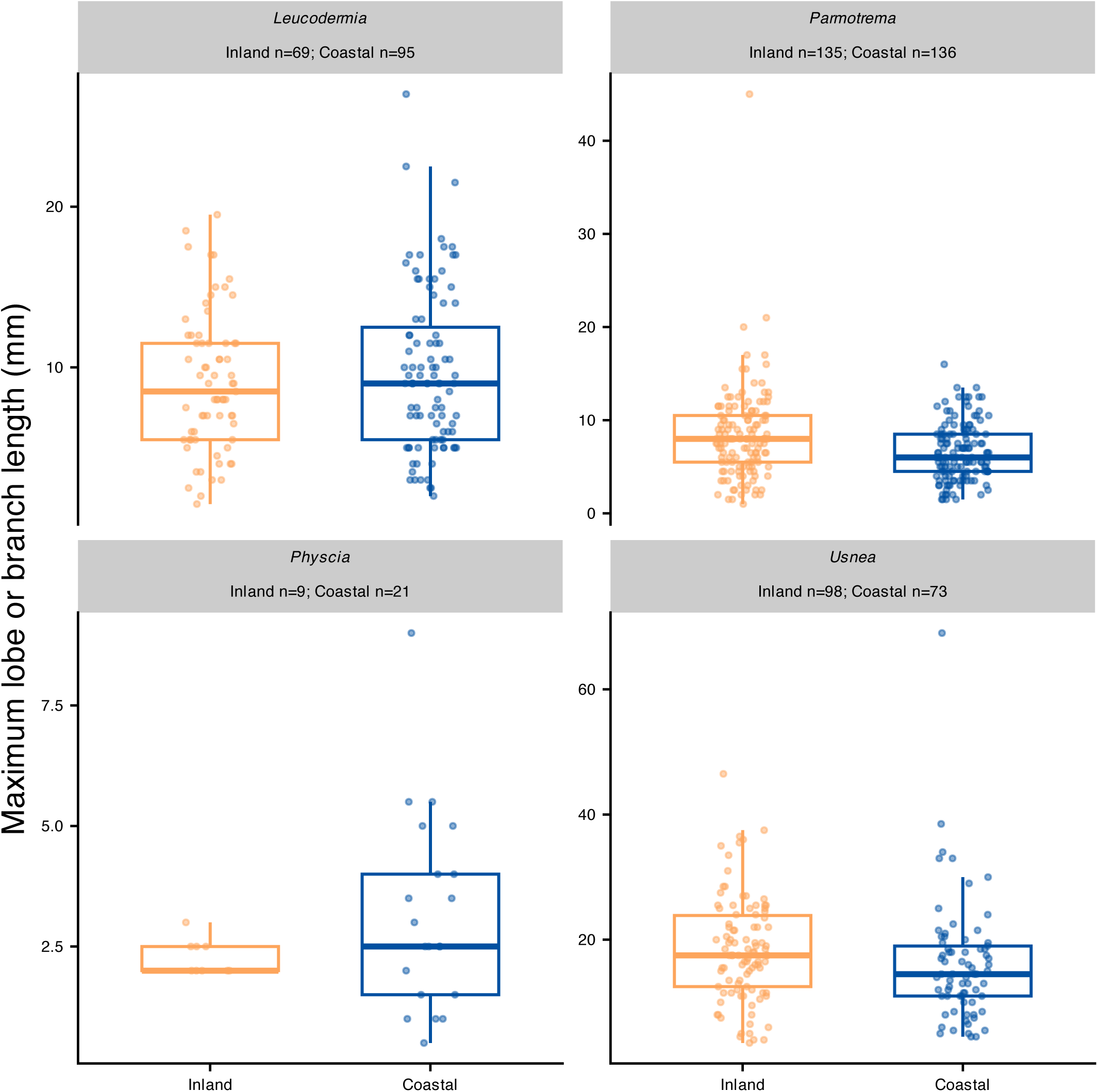
Taxon-specific sensitivity analysis of macrolichen maximum lobe or branch length. Genera are shown only when represented by at least five observations on each transect. Within-genus responses were heterogeneous, with no consistent inland–coastal direction across taxa. These exploratory distributions complement the genus-adjusted mixed-effects analysis reported in the main text.

**Figure S2.**
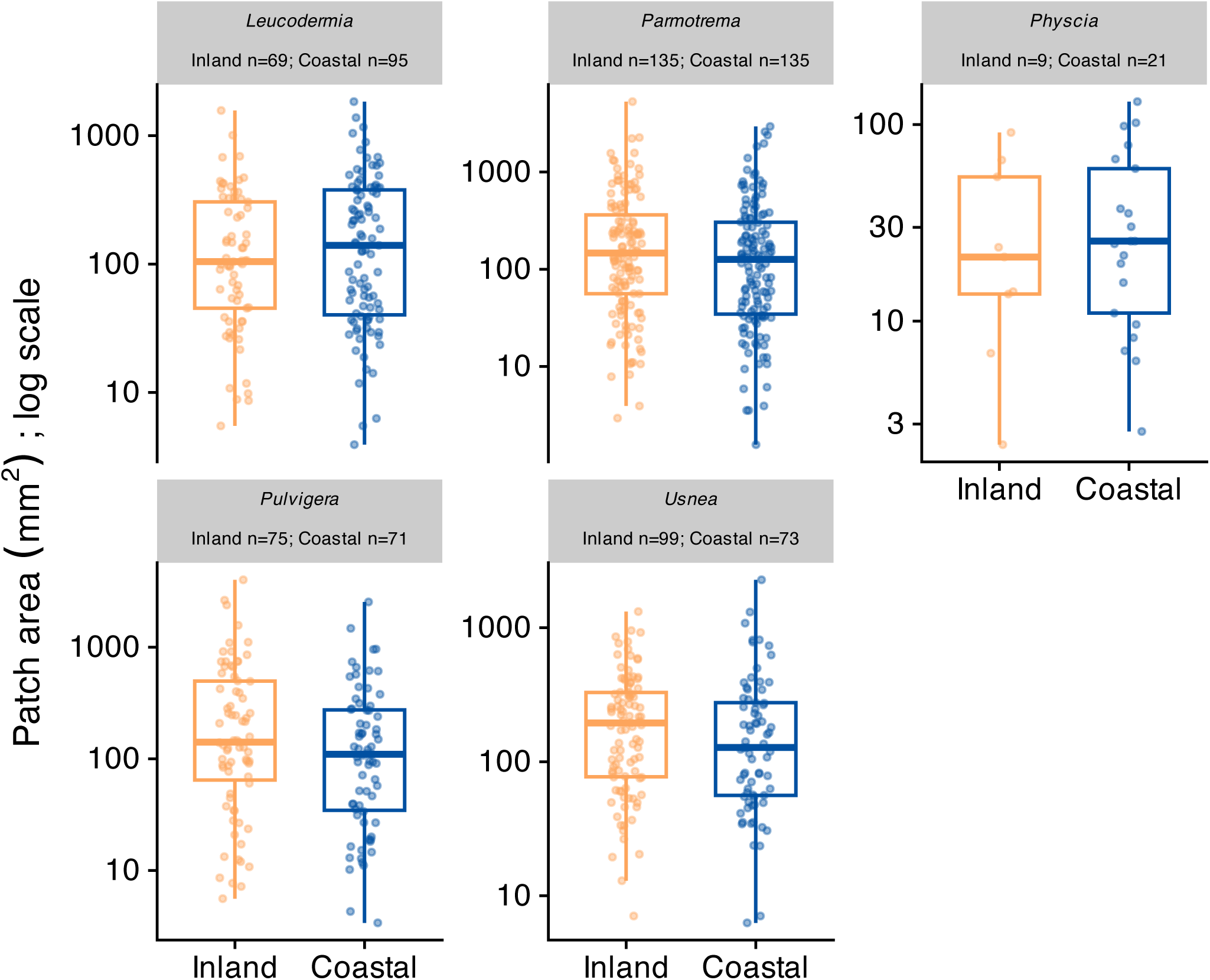
Taxon-specific sensitivity analysis of positive cryptogam patch area. Taxa are shown only when represented by at least five observations on each transect. Inland–coastal directions differed among taxa, indicating that the pooled patch-area difference did not reflect a uniform within-taxon response.

**Figure S3.**
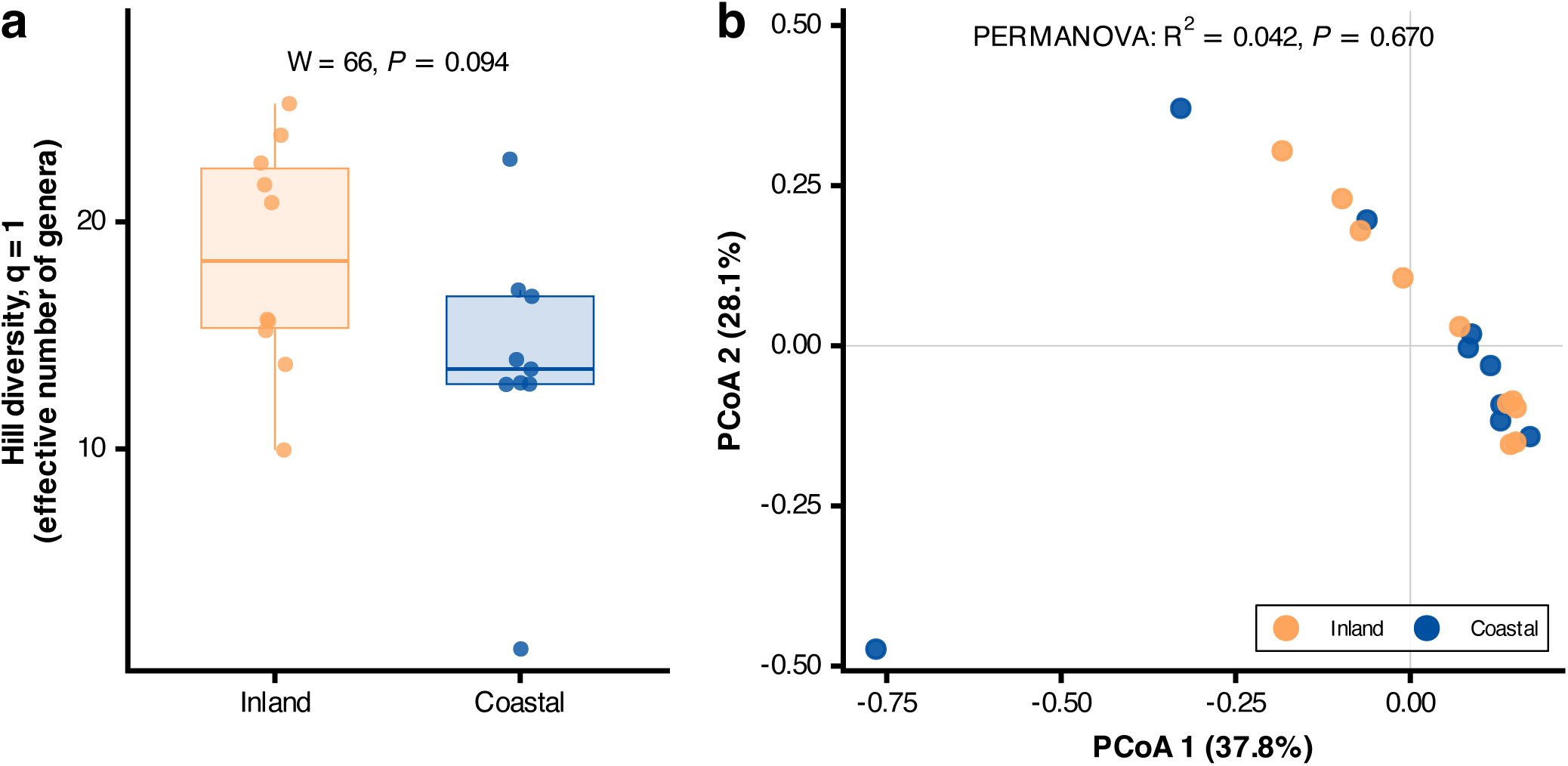
Bacterial community diversity and composition of Pulvigera pringlei across inland and coastal transects. (a) Genus-level Hill diversity (q = 1) of bacterial communities from inland and coastal samples. (b) Principal coordinates analysis of bacterial community composition based on Bray–Curtis dissimilarities; points are colored by transect. Multivariate dispersion did not differ between transects (permutation test: *P* = 0.469).

**Figure S4.**
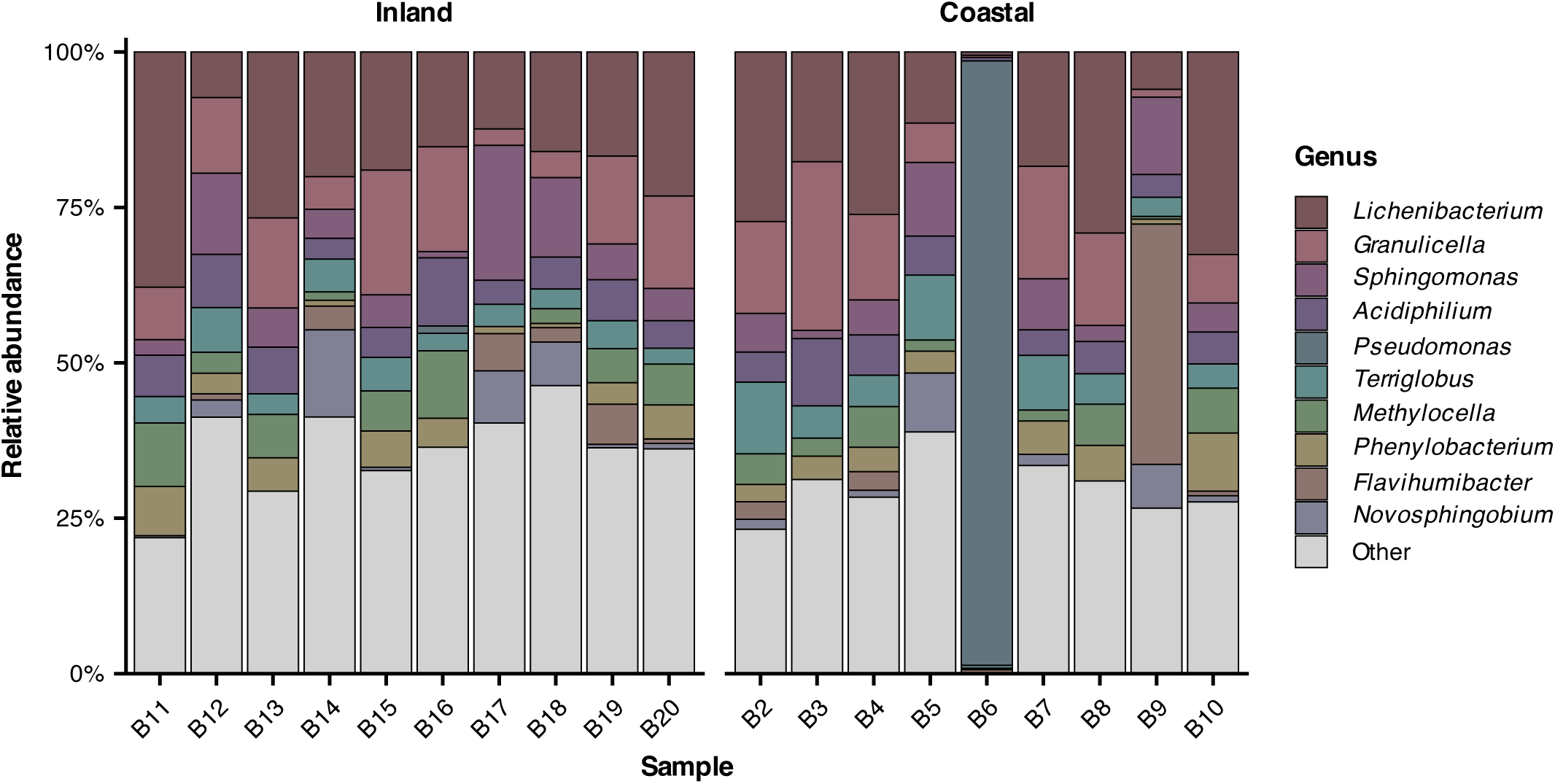
Bacterial community composition of Pulvigera pringlei. Relative abundance of the most common bacterial genera across inland and coastal *P. pringlei* samples.

**Figure S5.**
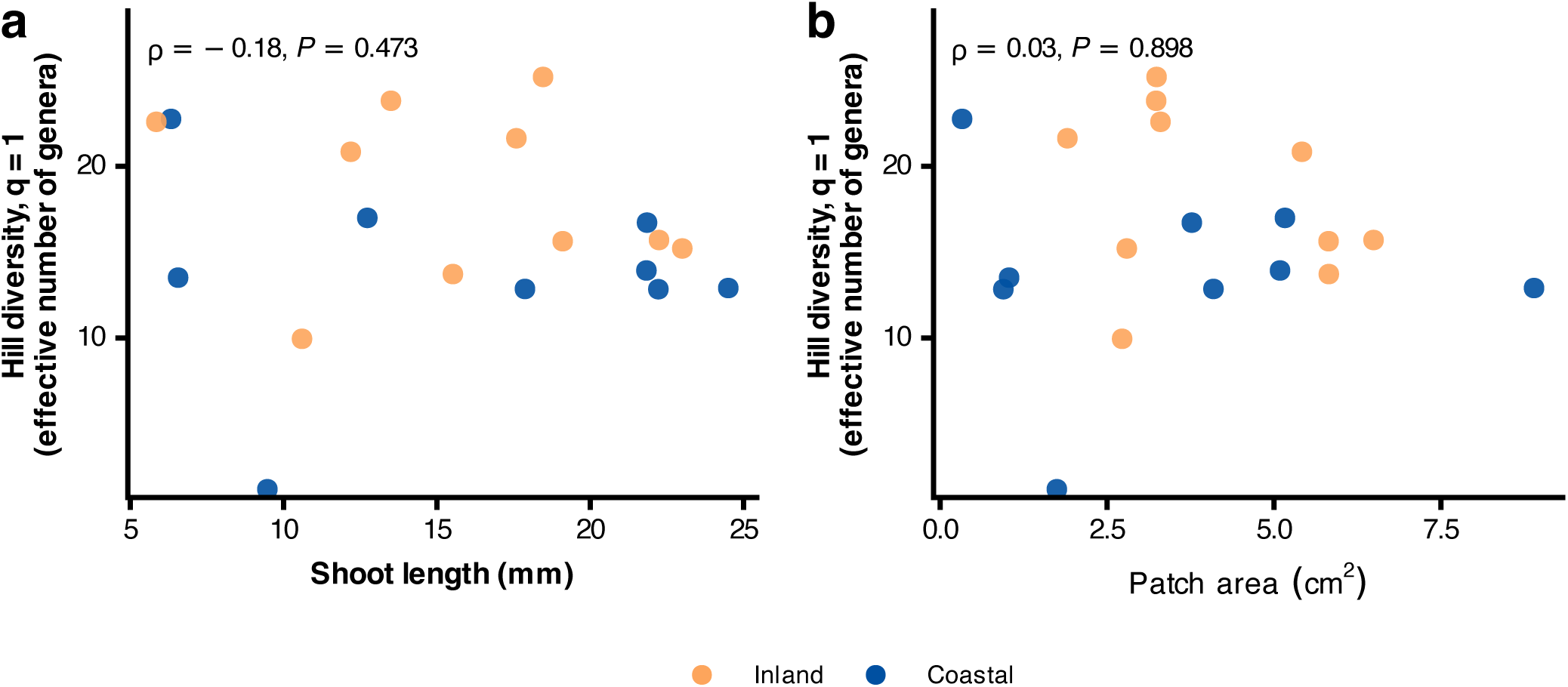
Associations between Pulvigera pringlei morphology and bacterial diversity. Relationships between bacterial genus-level Hill diversity (q = 1) and (a) moss shoot length and (b) moss patch area. Statistical tests were based on Spearman rank correlations.

## Supplementary Tables

**Table S1.**
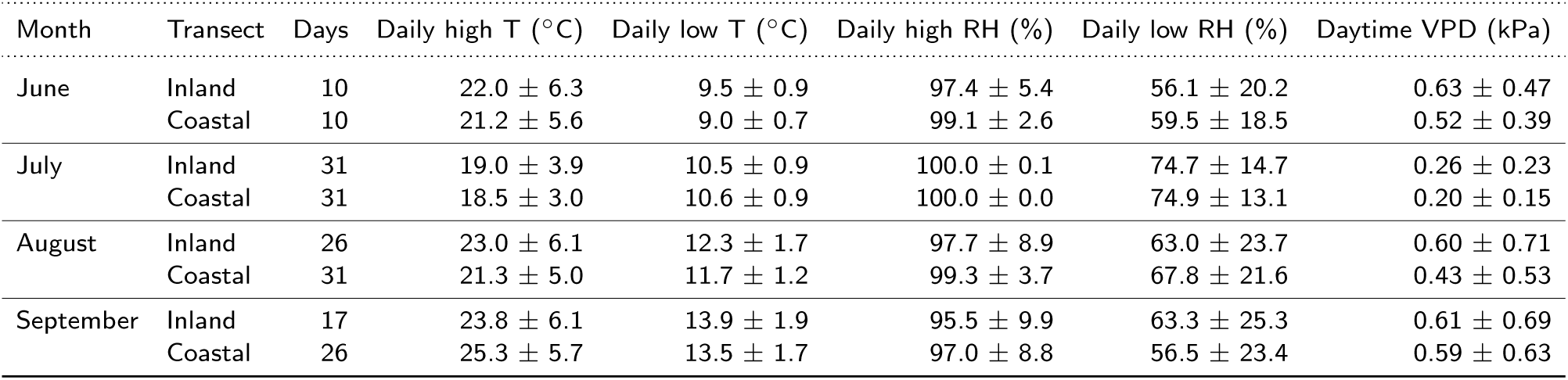
Monthly microclimate conditions on Inland and Coastal transects. Values are monthly mean *±* SD of daily conditions. Temperature and relative humidity summarize daily sensor-level maxima and minima; VPD is mean daytime vapor pressure deficit. Days indicates the number of dates contributing to each monthly summary.

**Table S2:**
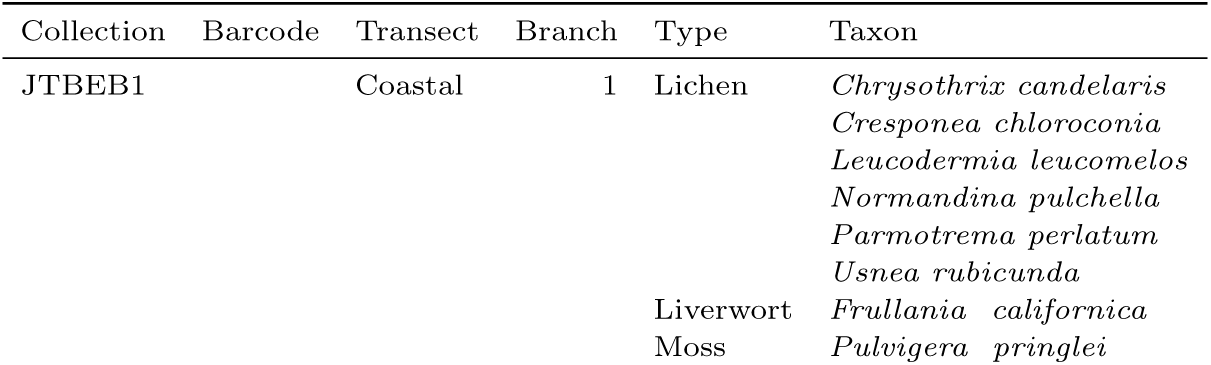
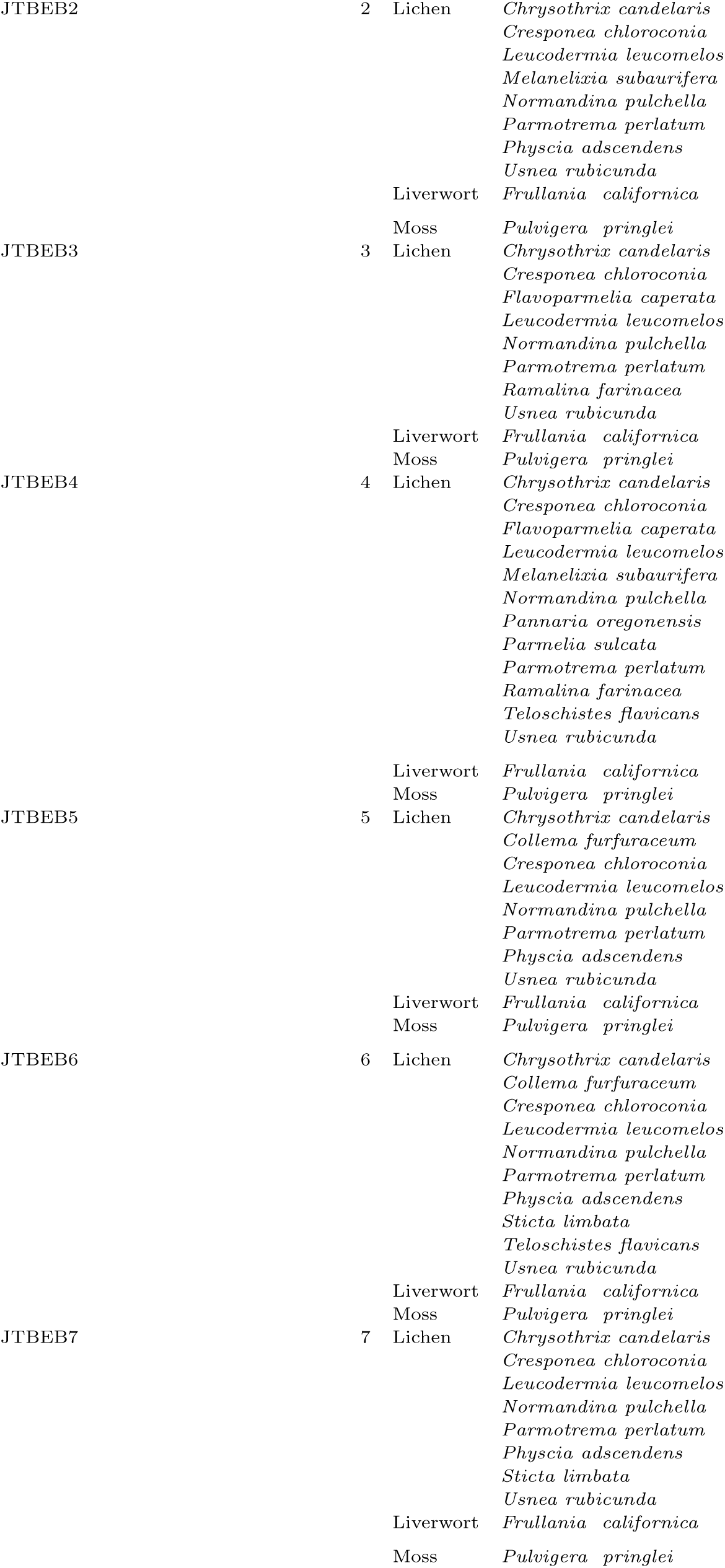
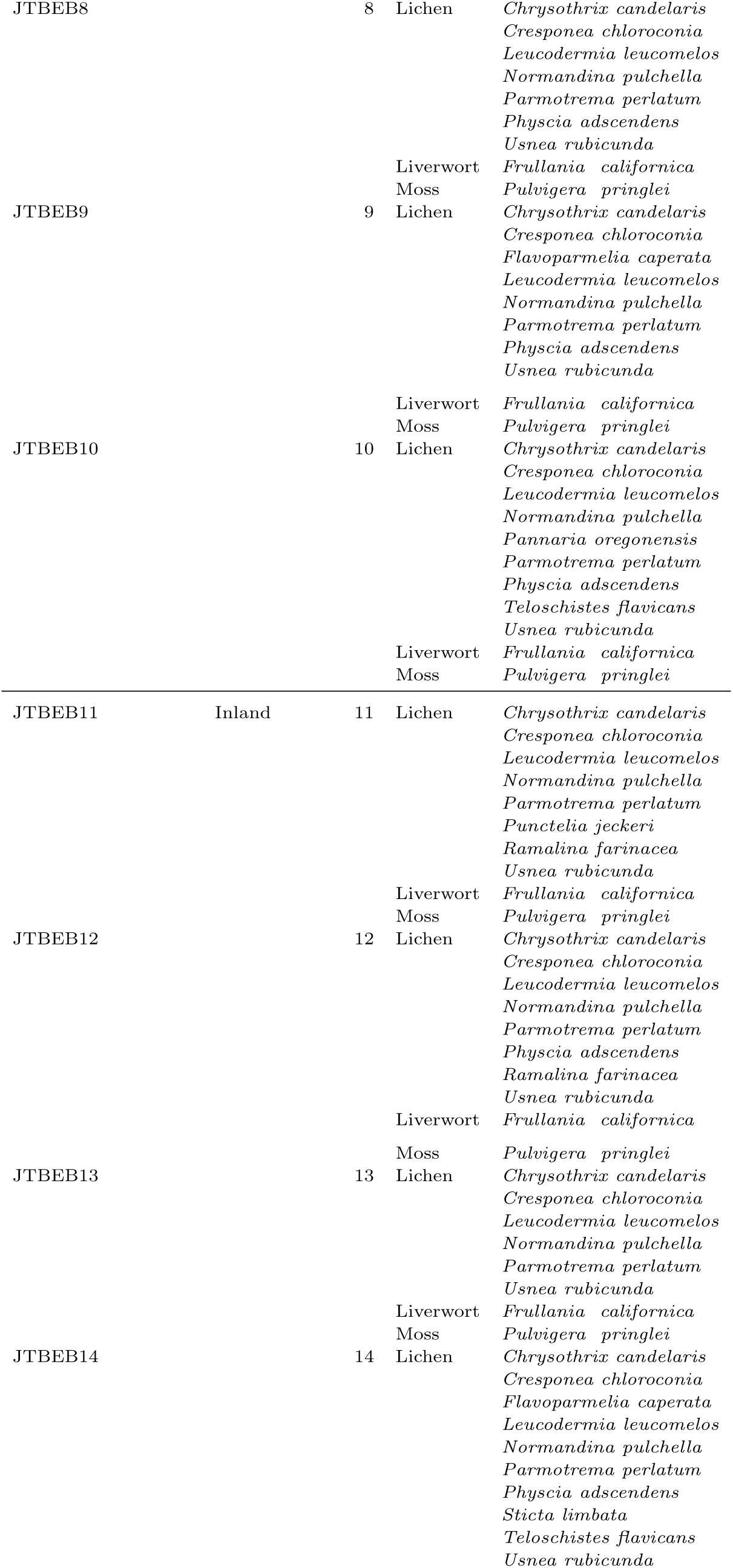
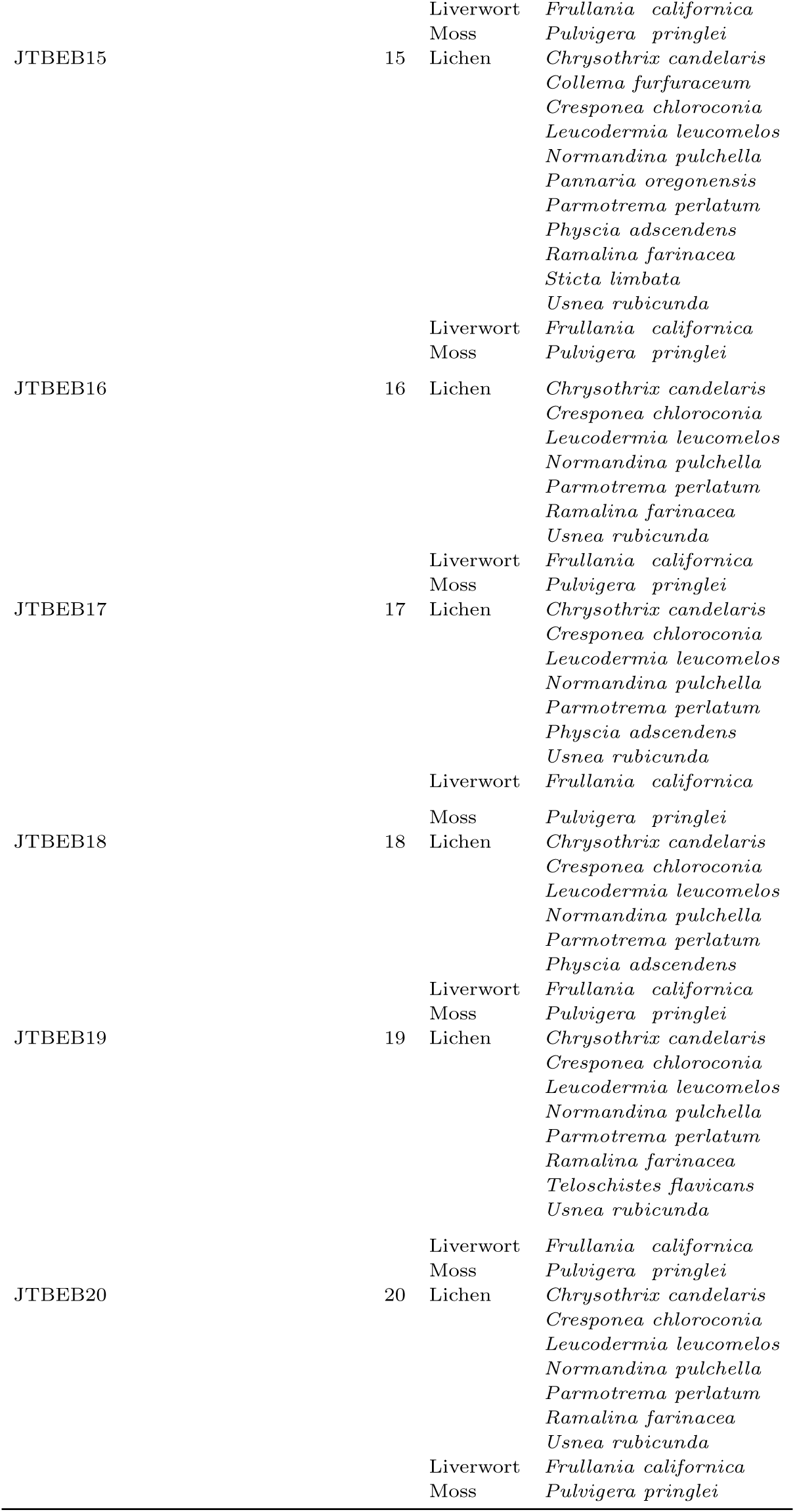
Herbarium voucher specimens representing cryptogam communities on sampled branches. Each collection represents a mixed-community voucher; taxa are listed by cryptogam group. Barcode fields are pending herbarium accession.

**Table S3.**
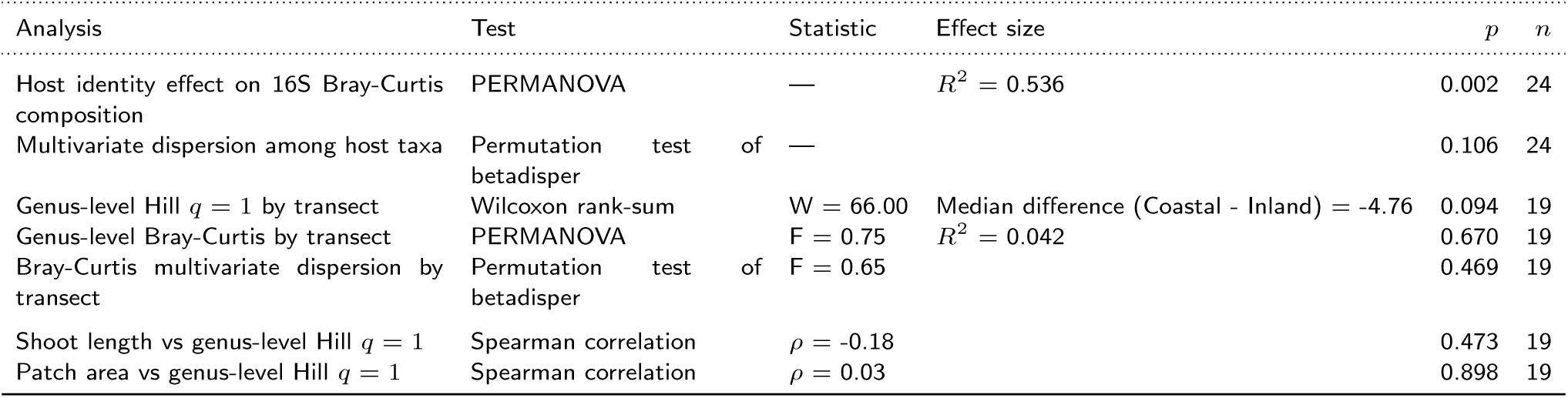
Summary of microbiome statistical analyses. For the genus-level Hill q = 1 comparison, the median difference is defined as Coastal minus Inland; negative values indicate a lower Coastal median.

**Table S4.**
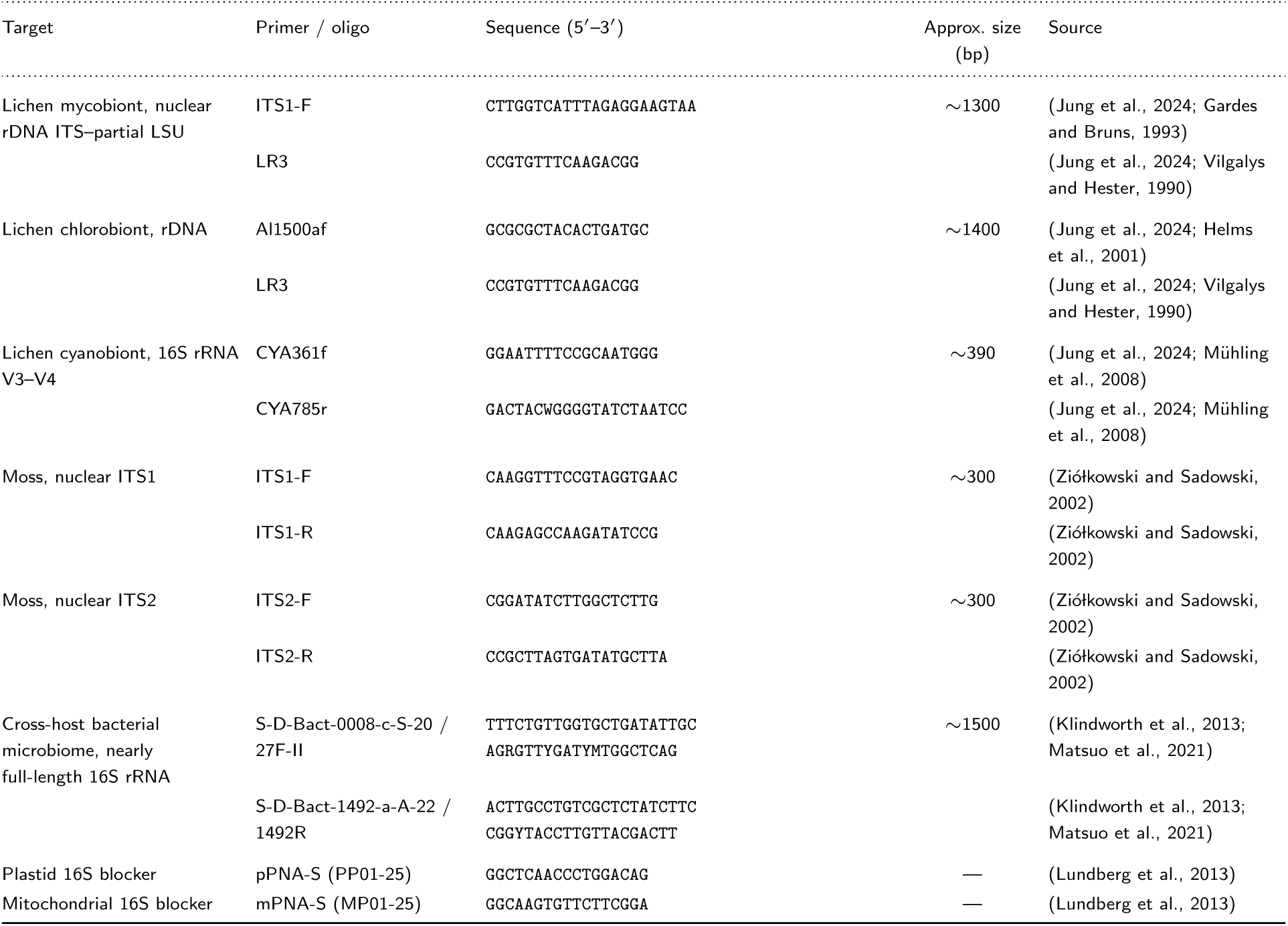
Primers and peptide nucleic acid (PNA) blockers used for cryptogam, symbiont, and cross-host bacterial 16S amplification. Amplicon sizes are approximate. Full-length bacterial 16S oligonucleotides include Oxford Nanopore-compatible 5*^′^* anchor sequences. The dedicated *Pulvigera pringlei* microbiome samples were amplified and sequenced separately by Plasmidsaurus and are not represented here.

